# MAPK Pathway Inhibition Reshapes Kinase Chemical Probe Reactivity Reflecting Cellular Activation States

**DOI:** 10.1101/2025.10.22.684065

**Authors:** Andrew F. Jarvis, Mohd Younis Bhat, Timothe Maujean, Ahlenne L. Abreu, Kaitlyn Toy, George M. Burslem, Donita C. Brady

## Abstract

Despite the pivotal role of oncogenic kinases in cancer initiation, progression, and therapeutic resistance, functionally profiling their activity and conformational dynamics in live cells remains challenging. Existing methods often fail to capture inhibitor-bound structural states of kinases, particularly in clinically relevant contexts such as treatment response and acquired resistance, where genomic data alone are insufficient. Here, we use activity-based protein profiling (ABPP) to monitor composite amino acid reactivity changes - across cysteine, lysine, and carboxylic acid residues - as a functional readout of kinase conformation and activation state. Using electrophilic probes, we show that treatment of BRAF^V600E^-mutant melanoma cells with vemurafenib or trametinib decreases overall cysteine and lysine reactivity in BRAF^V600E^ and MEK1/2, likely reflecting composite changes in amino acid accessibility across multiple reactive residues associated with inhibitor binding. Changing the order of probe addition and inhibitor treatment altered labeling outcomes, consistent with competitive engagement and structural stabilization. Comparative analysis of ATP-competitive BRAF^V600E^ inhibitors vemurafenib and dabrafenib revealed distinct impacts on aspartate and glutamate labeling patterns, suggesting that ABPP can distinguish inhibitor-dependent differences in residue accessibility that may reflect distinct inhibitor-bound conformations. In inhibitor-resistant melanoma models, ABPP detected differential residue reactivity relative to parental cells, consistent with BRAF overexpression and the MEK2 Q60P activation mutation, both established mechanisms of MAPK inhibitor resistance. Moreover, global proteome analyses of cysteine and lysine reactivity upon BRAF^V600E^ inhibition, revealed probe-accessible cysteine labeling changes in labeling on KSR2, indicating broader MAPK pathway remodeling. Together, these findings establish ABPP as a powerful chemical biology approach for investigating inhibitor-dependent changes in kinase residue accessibility, providing a framework to explore how conformational dynamics and pathway adaptation shape therapeutic response and resistance in oncogenic signaling networks.

## Introduction

Despite the central role of oncogenic proteins and their downstream effectors in driving tumor biology, integration of protein biochemistry into clinical oncology remains largely limited compared to genomic approaches reporting on mutation state. A key barrier is the biochemical specificity of each protein, requiring bespoke methods to assess functional states such as enzymatic activity, cofactor binding, and post-translational modifications (PTMs). Traditional protein assays often suffer from low throughput, limited multiplexing, and subjective interpretation, reducing clinical utility [1,2]. Moreover, many functionally relevant PTMs, cofactors, and their regulatory enzymes remain poorly annotated. To address these limitations, there is a critical need for proteome-centered platforms capable of directly profiling tumor-specific protein activities. Encouragingly, a number of emerging technologies—such as reverse phase protein arrays (RPPA), chemoproteomics, and multiplexed inhibitor bead mass spectrometry—are increasingly being adopted for quantitative assessment of proteomic alterations in tumors [3-5]. These approaches are now being applied to predict patient responses to standard therapies and, in some cases, to inform the development of novel therapeutic agents.

Activity-Based Protein Profiling (ABPP) is an established chemical biology technique that utilizes small-molecule probes featuring an electrophilic reactive group designed to selectively react with nucleophilic amino acid sidechains within target proteins. When integrated with mass spectrometry, ABPP has enabled substantial advances in proteome-wide target identification and assessment of protein reactivity across diverse biological contexts [6-10]. Originally developed to target reactive serine and cysteine residues, ABPP has since evolved to enable profiling of a broader range of amino acids, including commonly post-translationally modified residues such as lysine, tyrosine, aspartate, glutamate, and histidine, as well as less frequently modified—but biologically significant—residues like methionine and arginine [11-15]. This approach has revealed previously inaccessible insights into protein targets and their biological functions, while also establishing a new framework for covalent drug discovery. Notably, the iterative optimization of chemical probes to selectively engage reactive, functionally essential cysteine residues in oncogenic proteins has directly contributed to the development of clinically approved covalent inhibitors—most prominently those targeting KRAS G12C and mutant EGFR [16,17]. Beyond oncogene targeting, valuable community resources such as CysDB and DrugMap have systematically expanded our understanding of cysteine reactivity: CysDB through comprehensive profiling of cysteine ligandability across thousands of reactive sites, and DrugMap through large-scale analysis of druggable cysteine sites across 416 cancer cell lines [18,19]. These datasets not only inform on protein druggability but also reflect cellular states, including response to altered cellular metabolism or pharmacological perturbation with covalent inhibitors [18,19]. Notably, although a large and growing body of literature on activity-based protein profiling, including studies demonstrating competition between covalent drugs and amino acid-reactive probes [8], the use of ABPP to evaluate residue-level competition between widely used noncovalent inhibitors and electrophilic alkyne probes remains largely unexplored. This represents a key conceptual advance, as it extends ABPP beyond traditional covalent competition paradigms to evaluate how noncovalent inhibitor engagement alters residue reactivity in live cells.

Protein kinases are among the most druggable enzyme class, particularly in the context of precision oncology [20]. However, delineating kinase activation states and inhibitor engagement remains challenging. Current approaches such as RPPA, depend on the availability and specificity of both phospho-specific and total protein antibodies, and multiplexed inhibitor bead mass spectrometry (MIB-MS), which can detect over 380 kinases, provide useful activity profiles but does not directly report on kinase structural states or inhibitor-bound conformations [3-5]. As a result, much of our understanding of kinase conformational dynamics has been derived from *in vitro* biochemical assays using purified proteins and biophysical techniques such as X-ray crystallography, cryo-electron microscopy, and NMR spectroscopy. While invaluable, these methods offer only static snapshots of individual states, often divorced from their native physiological context. For example, despite extensive efforts, the active conformation of the MAP2K kinases MEK1 and MEK2 has proven difficult to capture crystallographically [21]. Historically, efforts to assess residue reactivity in protein kinases have primarily focused on cysteine reactivity, as cysteine residues often contribute to structural stabilization and cofactor binding across diverse kinases [22,23], making them attractive targets for covalent probe development. However, residues such as lysine—central to ATP coordination—and the DFG aspartate—required for catalytic activation—also present compelling opportunities for functional profiling. Supporting this expanded perspective, Liu *et* al. [24] used structural modeling to predict ligandable cysteine and lysine residues across the kinome, reinforcing a shift in the field toward broader reactivity mapping [25].

The conformational state of a kinase is a critical determinant of inhibitor binding. Type I inhibitors target the active conformation of the ATP-binding site, whereas Type II inhibitors preferentially bind the inactive form [21]. This conformational selectivity provides an opportunity to probe differential residue accessibility *in situ*, particularly at catalytically or structurally important residues such as lysine and aspartate, which have been hypothesized to change accessibility upon activation or inhibitor binding. Our understanding of kinase conformational regulation has also expanded within the identification of intermediate states, such as the DFG-inter conformation, and the emergence of Type 1 ½ inhibitors, which engage conformations distinct from classical active or inactive states [26,27]. These insights underscore the potential for amino acid-reactive chemical probes to serve as functional reporters of kinase conformational state by measuring localized changes in residue accessibility.

Leveraging advances in ABPP and chemoproteomics, we designed a series of chemical biology and proteomic experiments to visualize and quantitatively assess kinase amino acid reactivity changes in native cellular contexts following targeted inhibitor engagement, addressing a longstanding challenge in functional kinase profiling. Specifically, we aimed to overcome persistent barriers in distinguishing active and inactive kinase states by using amino acid-reactive probe labeling as a proxy for conformational state and inhibitor occupancy, thereby enabling functional insights into kinase regulation *in situ*. Furthermore, we hypothesized that proteome-wide reactivity in response to targeted kinase inhibition would potentially reveal molecular adaptations to therapy, including reactivity signatures associated with altered pathway signaling. An informative model system for evaluating such reactivity changes is BRAF^V600E^-driven melanoma, where hyperactivation of the MAPK signaling cascade drives tumor progression and metastasis. The V600E mutation in BRAF occurs in approximately 50% of melanoma cases [28], has therefore been a major focus of therapeutic development. Multiple FDA-approved inhibitors target BRAF^V600E^ and its downstream effectors, such as MEK1/2, each engaging distinct binding modes and conformational states. For example, vemurafenib binds the active conformation of BRAF^V600E^ at the ATP-binding site, whereas trametinib targets binds an allosteric pocket in in MEK1/2 specific to the inactive state [29-32]. Despite these advances, clinical response rates to MAPK pathway inhibitors remain limited, with only 10–15% of patients achieving durable benefit [33]. While resistance mechanisms, including BRAF amplification and the MEK2 Q60P activating mutation, have been identified, their structural and biochemical consequences remain poorly defined [34]. This underscores the need for approaches that probe kinase structure and reactivity directly within cells, particularly in drug-resistant contexts, to enable the development of next-generation inhibitors informed by functional data.

Here, we expand the conceptual and methodological scope of ABPP by integrating composite amino acid reactivity profiling, measuring reactivity across cysteine, lysine, aspartate, and glutamate residues, to infer changes in protein conformation and activation state. This approach enables visualization of inhibitor-associated alterations in residue accessibility that likely correspond to changes in kinase structural states *in situ*, bridging the gap between biochemical reactivity and conformational dynamics within live cells. Using this strategy, we investigated BRAF^V600E^ and MEK1/2 reactivity in response to kinase inhibition and acquired drug resistance, comparing residue accessibility following treatment with inhibitors that engage distinct conformations and binding sites. In parallel, we conducted proteome-wide profiling to capture broader reactivity shifts across the MAPK pathway, revealing node-specific responses dependent on the point of inhibitor engagement. To assess clinical relevance, we examined melanoma cell lines with acquired resistance to BRAF^V600E^ or MEK1/2 inhibitors, uncovering resistance-associated alterations in kinase reactivity consistent with BRAF amplification and the MEK2 Q60P activating mutation. Together, these findings establish ABPP as a powerful chemoproteomic platform for interrogating kinase conformational states, inhibitor binding, and resistance mechanisms *in situ*, linking local amino acid reactivity to structural dynamics and therapeutic adaptation within oncogenic signaling networks.

## Results

### MEK1/2 and BRAF Inhibitors Suppress Overall Cysteine and Lysine Accessibility in Target Proteins

We hypothesized that MAPK pathway inhibitors alter the overall chemical accessibility of reactive amino acids within their target kinases, reflecting changes in composite residue reactivity associated with inhibitor engagement. To test this, we focused on two well-characterized, FDA-approved MAPK inhibitors with distinct binding modes: the ATP-competitive BRAF inhibitor vemurafenib and the allosteric MEK1/2 inhibitor trametinib. Using BRAF^V600E^-mutant A375 melanoma cells as a model system, we asked whether short-term treatment with these inhibitors produces measurable shifts in cysteine and lysine probe labeling that correspond to altered accessibility in BRAF^V600E^ and MEK1/2 (Figure 1). Cells were treated for one hour with 1 μM vemurafenib, 100 nM trametinib, or vehicle control, followed by one-hour in situ labeling with amino acid–selective activity-based probes: 25 μM iodoacetamide-alkyne (IA-alkyne) to assess cysteine reactivity and 25 μM ArSq-alkyne to assess lysine reactivity (Figure 2A, 2B, and S1, n =3). Then, the samples underwent a copper catalyzed click reaction with a TAMRA-biotin azide for visualization of changes through in-gel fluorescence, or with a biotin azide for streptavidin pulldown, which was followed by both immunoblotting and quantitative LC-MS (Figure 2A). This experimental design was intended to determine whether inhibitor binding modifies probe accessibility across multiple reactive residues within target kinases, providing a functional, composite readout of inhibitor engagement in living cells. Initial in-gel fluorescence confirmed robust probe reactivity across the proteome and revealed visible global labeling differences between inhibitor-treated and control samples (Figure S2). As expected, immunoblot analysis verified that both inhibitors reduced MEK1/2 and ERK1/2 phosphorylation (Figure 2C, Figure S3A and S3B, n = 3), providing indirect evidence for target engagement, with no impact observed in probe only conditions. Following click chemistry and streptavidin enrichment, we observed reduced recovery of endogenous MEK1/2 and BRAF^V600E^ in the presence of their respective inhibitors compared to probe-only conditions (Figure 2C, Figure S3A and S3B, n = 3). These decreases reflect reduced composite cysteine and lysine reactivity, consistent with inhibitor binding altering overall residue accessibility within each kinase. Quantitative densitometry across three replicates revealed significant decreases in probe-enriched MEK1/2 after trametinib treatment (IA-alkyne: 0.35 ± 0.05; ArSq-alkyne: 0.37 ± 0.04, p < 0.001) and in BRAF^V600E^ after vemurafenib treatment (IA-alkyne: 0.24 ± 0.07; ArSq-alkyne: 0.41 ± 0.08, p < 0.05) (Figure 2E and 2F). Reversing the order of treatment—adding probes before inhibitors—abolished these effects (Figure 2D, Figure S4A–C, n = 3), indicating that the observed decreases depend on prior inhibitor engagement rather than nonspecific labeling artifacts.

**Figure 1.**
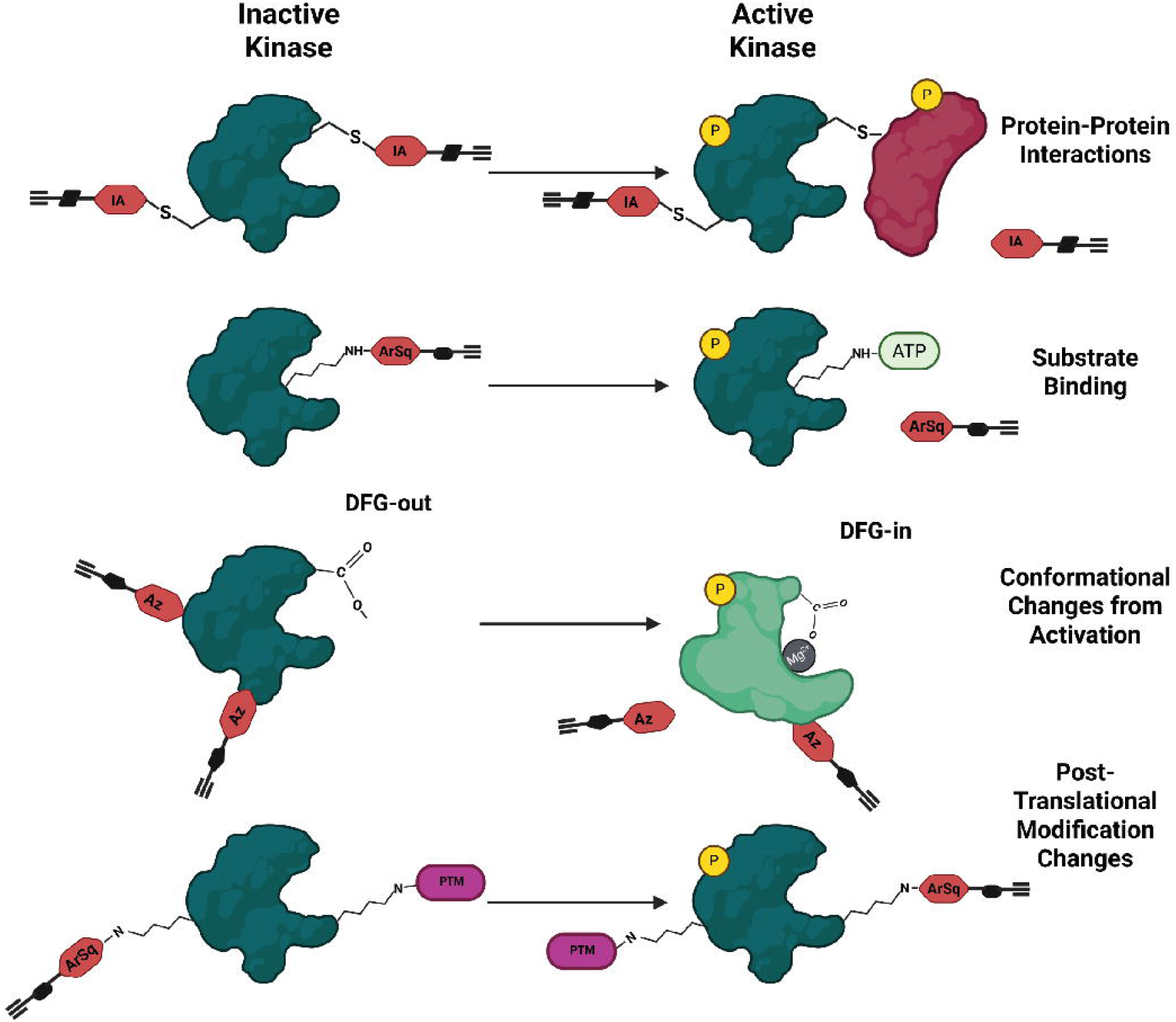
Activity Based Protein Profiling Can Use Broad Residue Accessibility Changes to Drive Hypotheses about Protein Activity in Live Cells. From readouts in changes in residue accessibility of cysteine, lysine, aspartate, and glutamate, Activity Based Protein Profiling can provide information in light of activity changes of Kinases, such as alterations in protein-protein interactions, substrate binding, conformational changes, and/or post-translational modifications.

**Figure 2.**
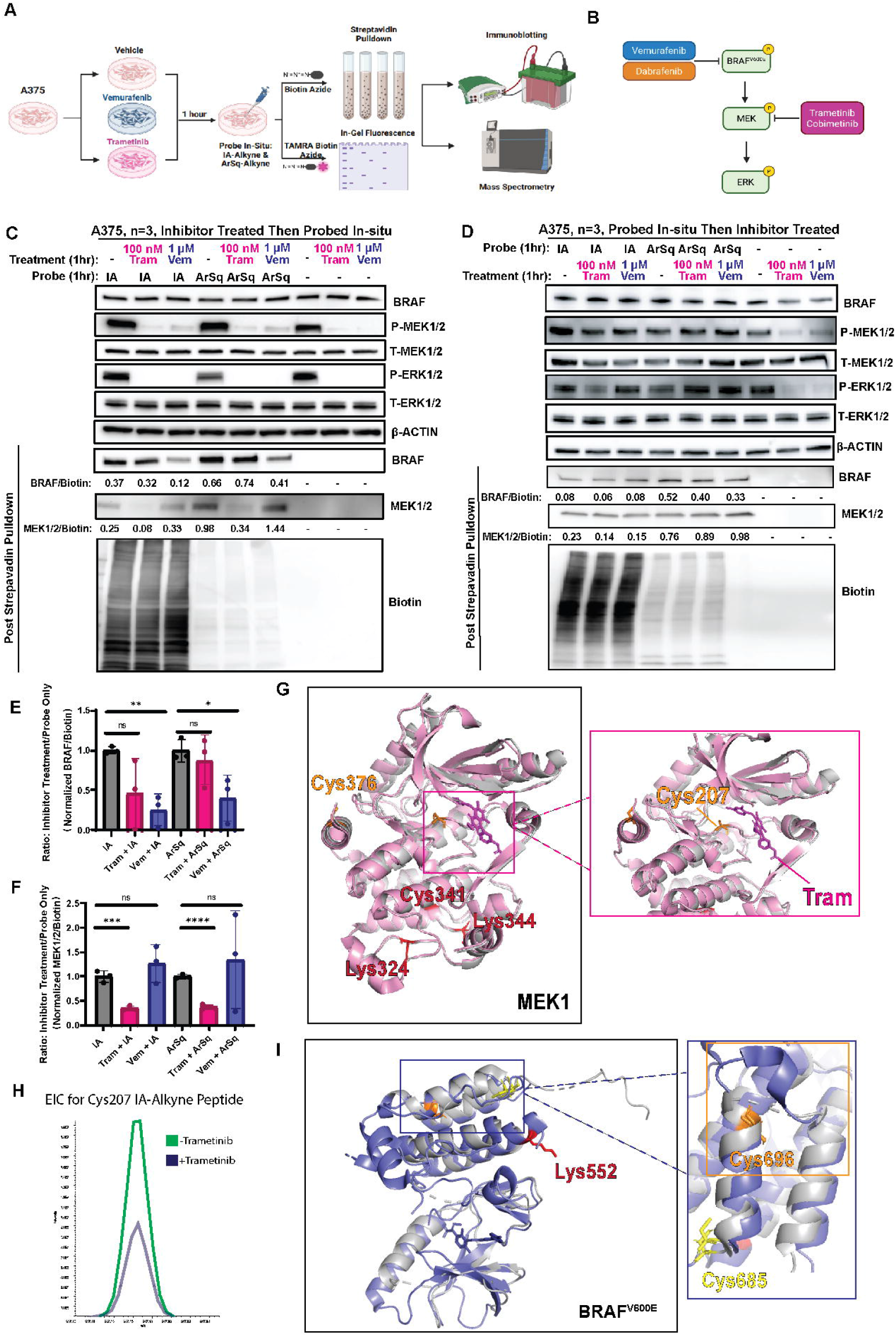
Targeted inhibition of BRAF^V600E^ and MEK1/2 reduces their cysteine and lysine residue accessibility. (A) Schematic overview of experimental workflow for assessing residue accessibility in BRAF^V600E^-mutant melanoma cells. (B) Diagram illustrates the oncogenic MAPK signaling cascade initiated by BRAFV600E and highlights FDA-approved inhibitors targeting mutant BRAF and its downstream effectors MEK1/2. This pathway plays a critical role in melanoma progression and is a key target for therapeutic intervention. (C) Top: Immunoblot detection of phosphorylated (P)-MEK1/2, total (T)-MEK1/2, P-ERK1/2, T-ERK1/2, and β-actin from A375 cells treated for one hour with vehicle, 100 nM trametinib, or 1μM Vemurafenib, and labeled *in situ* for one hour with no probe, IA-alkyne, or ArSq-alkyne.Bottom: Immunoblot detection of BRAF, MEK1/2, or biotin after streptavidin pull-down from A375 cells treated with vehicle, 100 nM Trametinib, or 1μM Vemurafenib, for 24 hours and labeled with no probe, 25μM IA-alkyne, or 25μM ArSq-alkyne followed by Cu-catalyzed click chemistry with biotin azide. (D) Immunoblot detection of phosphorylated (P)-MEK1/2, total (T)-MEK1/2, P-ERK1/2, T-ERK1/2, and β-actin from A375 cells and immunoblot detection of BRAF, MEK1/2, or biotin after streptavidin pull-down from A375 cells first treated with no probe, IA-alkyne, or ArSq-alkyne for one hour and then with treated with vehicle, 100 nM Trametinib, or 1μM Vemurafenib for one hour followed by Cu-catalyzed click chemistry with biotin azide. (E) Densitometry quantification of three replicates of the streptavidin pulldown of MEK as in Figure 2C. MEK band intensities normalized to whole-lane Biotin and then to vehicle control. Results were compared using a pair-wise t-tests. One asterisk, P<0.05, Two asterisks, P<0.01, Three asterisks, P<0.001, Four asterisks, P<0.0001. n=3. (F) Densitometry quantification of three replicates of the streptavidin pulldown of BRAF as in Figure 2C. BRAF band intensities normalized to whole-lane Biotin and then to vehicle control. Results were compared using a pair-wise t-tests. One asterisk, P<0.05, Two asterisks, P<0.01, Three asterisks, P<0.001, Four asterisks, P<0.0001. n=3. (G) MEK1 unbound structure overlayed with trametinib bound MEK1, Sites of IA-alkyne and ArSq-alkyne labeling in MEK1 either after enrichment of overexpressed Flag-MEK1 (yellow) or from purified protein (red), or both (orange). (H) Extracted Ion Chromatogram from MEK1 C207 labeling with IA-alkyne. (I) BRAF V600E unbound (gray) overlayed with BRAF^V600E^ Vemurafenib bound, Identified Sites of IA-alkyne, ArSq-alkyne and Az-alkyne Labeling in BRAF^V600E^ either after enrichment of overexpressed Flag-BRAFV600E (yellow) or from purified protein (red), or both (orange), with zoom in on Cys696 and surrounding structures with difference highlighted in box.

To verify that these differences arose from inhibitor competition rather than abundance changes, we performed parallel lysate labeling after 24-hour inhibitor treatment. Comparable patterns of probe reactivity were observed in vitro and in situ (Figure S5A–C, n = 3), confirming that the measured changes represent altered probe accessibility rather than protein loss. Collectively, these experiments establish that vemurafenib and trametinib each reduce the overall cysteine and lysine reactivity of their cognate targets under conditions consistent with direct inhibitor engagement. Curiously, longer treatment with the cysteine- or lysine-targeted activity-based probes led to a similar decrease in MEK1/2 and ERK1/2 phosphorylation, along with reduced cystine and lysine reactivity (Figure S6A and S6B, n=3). To focus on the acute impacts of kinase inhibition, we focused on one hour treatment experiments, followed by one hour of *in situ* probe labelling for the rest of this study.

To pinpoint reactive residues that contribute to these composite changes, we pursued orthogonal approaches to identify reactive cysteine and lysine residues within BRAF^V600E^ and MEK1/2. First, we mapped probe labeling sites in MEK1 using recombinant protein and FLAG-tagged MEK1 overexpression in A375 cells. Biochemical labeling of MEK1 *in vitro* identified cysteine adducts at C207, C341, and C376, and lysine adducts at K324 and K344 (Figure 2G and Figure S7A). *In situ* FLAG-MEK1 labeling reproduced adducts at C207 and C376 (Figure 2G and Figure S7B). Trametinib treatment reduced labeling of MEK1 at C207 both *in situ* (Figure 2H) and biochemically (Figure S7C), which lies near the trametinib-binding pocket (Figure 2G) (PDB 7JUR) [29], supporting that local structural protection or steric competition limits probe access to multiple residues within the allosteric region. Although lysine labeling of MEK1 decreased upon trametinib treatment, specific reactive lysines could not be assigned *in situ*, indicating that the signal likely reflects changes across several lysines rather than one defined site.

We performed a similar cysteine and lysine site of labelling analysis with FLAG-BRAF^V600E^ overexpression in A375 cells (Figure 2I, Figure 3D) and recombinant kinase domain of BRAF^V600E^ (residues 416-766) (Figure 2I, Figure 3D, and Figure S8) and identified labelling at Cys 685, 696, 748, and Lys 253 and 552 both within the kinase domain and in regulatory domains of BRAF (Figure 2I, Figure 3D and Figure S8). While site-specific changes based on vemurafenib binding could not be unambiguously resolved, differential pulldown between inhibitor-treated and untreated samples is consistent with crystallographically observed differences between the apo and inhibitor-bound BRAF^V600E^ structures (Figure 2I). Together, these findings show that MAPK pathway inhibitors produce reproducible, protein-specific decreases in composite cysteine and lysine reactivity, consistent with inhibitor engagement reducing the accessibility of multiple reactive residues within the kinase active and allosteric regions.

**Figure 3.**
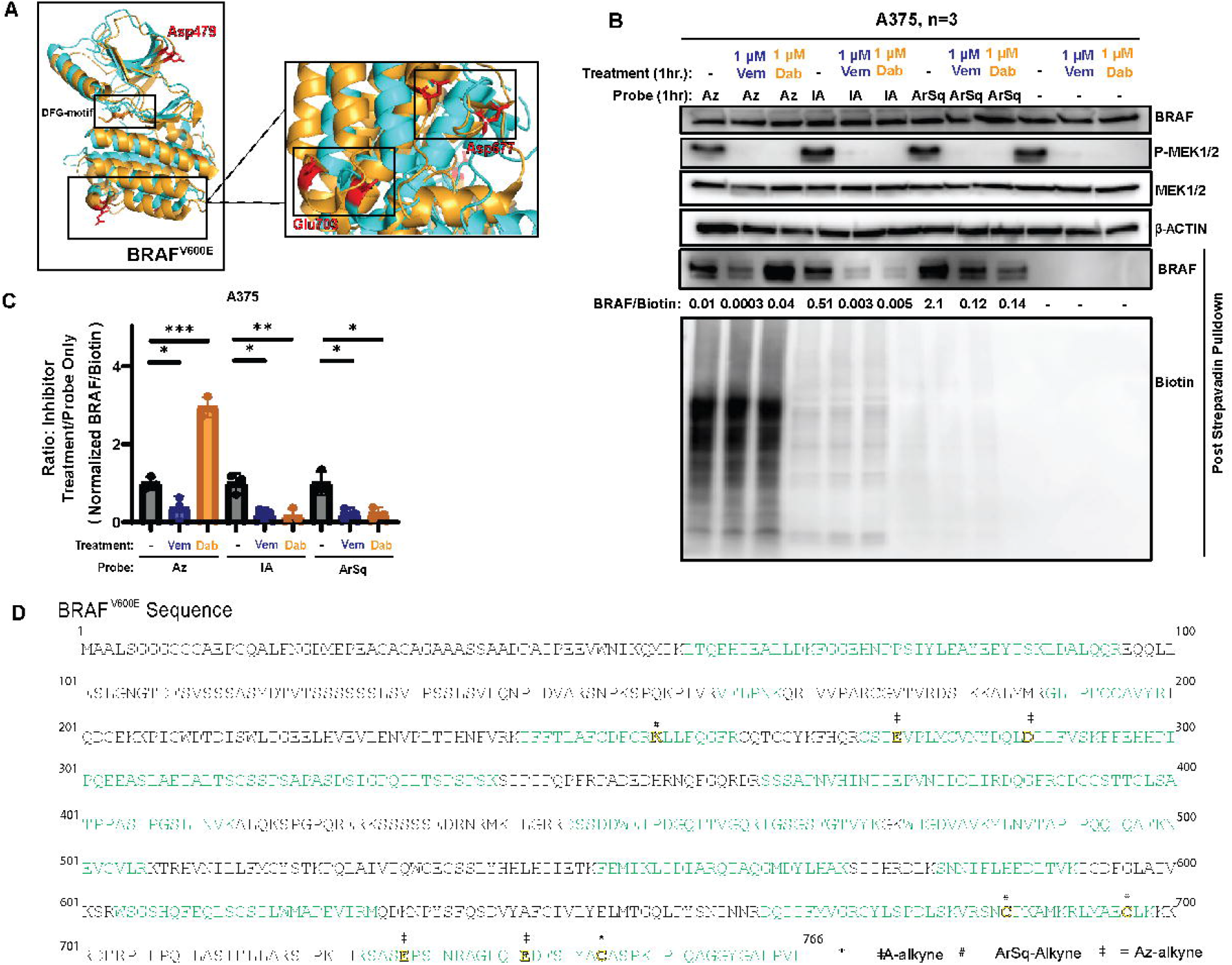
Vemurafenib and dabrafenib differentially alter carboxylic acid residue accessibility in BRAF. (A) Overlayed crystal structure of Dabrafenib bound to BRAF^V600E^ (orange) or Vemurafenib bound to BRAF^V600E^ (blue) with zoom in on Glu703 and Asp677 residue separation in each inhibitor bound conformation(B) Immunoblot detection of phosphorylated (P)-MEK1/2, total (T)-MEK1/2, and β-actin from A375 cells treated with vehicle, 1μM Vemurafenib, or 1μM dabrafenib for one hour and labeled for one hour with no probe, 10μM AZ-alkyne, 25μM IA-alkyne, or 25μM ArSq-alkyne. Immunoblot detection of BRAF or biotin after streptavidin pull-down from A375 cells treated with vehicle, 1μM Vemurafenib, or 1μM Dabrafenib, for one hour and labeled with no probe, AZ-alkyne, IA-alkyne, or ArSq-alkyne followed by Cu-catalyzed click chemistry with biotin azide. (C) Quantification: BRAF band intensity normalized to whole-lane Biotin and then to vehicle control. Results were compared using pair-wise t-tests. One asterisk, P<0.05, Two asterisks, P<0.01, Three asterisks, P<0.001, Four asterisks, P<0.0001. n=3. (D) Sequence Map for BRAF^V600E^ with annotated sites of labeling from IA-alkyne, ArSq-alkyne and Az-alkyne identified in overexpressed Flag-BRAF^V600E^.

### Distinct Patterns of Acidic Residue Reactivity Distinguish BRAF^V600E^ Inhibitor Classes

Having established that kinase reactivity can be assessed through changes in the accessibility of key functional residues, we next investigated whether different BRAF inhibitors, dabrafenib and vemurafenib, produce distinguishable effects on reactive acidic residues (Figure S2B and S6C). When overlaying structural models of BRAF^V600E^ bound to vemurafenib and dabrafenib (PDB ID: 4XV2), we observed that F595 adopts an outward, intermediate conformation in the dabrafenib-bound structure compared to its orientation in the vemurafenib-bound form (Figure 3A). This conformational shift also influenced the positioning of the adjacent D594 within the DFG loop. Using the aspartate and glutamate-selective probe Az-alkyne, we profiled residue accessibility under one-hour treatments with either inhibitor (Figure S1). Consistent with earlier observations, IA- and ArSq-alkyne labeling of BRAF^V600E^ decreased following both treatments (Figure 3B and 3C, Figure S9, n = 3). However, Az-alkyne labeling revealed divergent behavior: vemurafenib treatment reduced acidic-residue labeling (average fold change 0.43; p = 0.0337), whereas dabrafenib increased it nearly three-fold (p = 0.0004). These opposite effects suggest that the two inhibitors differentially influence the local environments of multiple acidic residues within BRAF^V600E^. This reduction coincided with the expected decrease in MEK1/2 phosphorylation, reflecting effective inhibition of oncogenic BRAF^V600E^ signaling (Figure 3B). Given the proximity of these residues to the kinase activation segment—particularly the DFG motif—these changes may reflect broader structural consequences of inhibitor binding that lead to the modulation of BRAF^V600E^ conformation, activity, or interaction states – all of which could be possible explanations for the differential probe labeling. To directly identify reactive acidic residues, we labeled both recombinant BRAF^V600E^ and overexpressed FLAG–BRAF^V600E^ with Az-alkyne. Mass spectrometry analysis revealed Glu 275, 730, 741 and Asp 287 in cellular samples (Figure 3D) and in Asp 479 and 677 and Glu 703 and 730 in the recombinant protein (Figure S8), as probe-reactive residues. Within the structural models of BRAF^V600E^ bound to vemurafenib and dabrafenib (PDB ID: 4XV2), we observed that the positions of D479, D677, and E703 that were labeled in Az-alkyne *in vitro* or *in situ* (Figure 3D and Figure S8) were distinct in the dabrafenib-bound structure compared to its orientation in the vemurafenib-bound form (Figure 3A). With only partial coverage of BRAF^V600E^ from tryptic digests (Figure 3D and Figure S8), it’s tempting to speculate but challenging to accurately determine the sites of labeling that determine this change. Future studies will focus on increasing our coverage of BRAF (especially the DFG loop) to determine if our hypothesis about Asp594 providing the predominant change in Az-Alkyne pulldown between dabrafenib or vemurafenib is correct (Figure 3A).

Applying the same approach to MEK1/2, we compared cysteine and lysine reactivity following treatment with the allosteric inhibitors trametinib and cobimetinib (Figure S10). Although both inhibitors bind near Cys 207 in MEK1, cobimetinib uniquely engages a region proximal to Lys 97 and exhibits a stronger preference for the active conformation of MEK1 (Figure S10A) [35]. Both drugs decreased MEK1/2 probe labeling to a similar extent (Figure S10B), indicating that composite reactivity changes in MEK1/2 are not solely determined by these inhibitor conformational preferences. Collectively, these results demonstrate that residue-selective probes can differentiate between inhibitor-specific effects on amino acid accessibility, revealing nuanced distinctions in how related inhibitors engage their targets in living cells.

### Drug-Resistant Melanoma Cells Exhibit Altered Composite Cysteine and Lysine Reactivity

To test whether MAPK inhibitor resistance alters kinase reactivity profiles (Figure S2C, S6D, and S6E), we analyzed parental and resistant 451-Lu melanoma cell lines. The BRAF inhibitor–resistant (BiR) cells overexpress BRAF (Figure 4A), while the MEK inhibitor–resistant (MiR) cells harbor the activating MEK2 Q60P mutation (Figure 4B). Both lines show sustained MEK1/2 and ERK1/2 phosphorylation despite inhibitor treatment (Figure 4C and 4E, Figure S6D, Figure S11, n = 3). Due to this aberrant MAPK pathway signaling, the inhibitor-resistant cells are normally grown in media containing either 1 μM vemurafenib (BRAFi-resistant) or 100 nM trametinib (MEKi-resistant) and require a four-hour washout to remove chronic drug exposure prior to treatment for with the appropriate inhibitor followed by in situ probe labeling (Figure 2A).This strategy also aimed to evaluate the potential of activity-based proteome profiling as a tool to monitor treatment response and resistance. In the parental line, vemurafenib and trametinib decreased cysteine and lysine labeling of BRAF^V600E^ and MEK1/2, mirroring A375 behavior (Figure 4C–F, Figure S11A and S11B). In contrast, resistant cells displayed increased probe reactivity for the corresponding kinase: IA-alkyne labeling of BRAF^V600E^ increased 1.5-to 1.9-fold in BiR cells, and MEK1/2 labeling increased two-fold in MiR cells (p < 0.05 for both) (Figure 4C-F, Figure S11A and S11B). These differences remained significant with or without inhibitor treatment. This increase in probe binding to MEK1/2 was also observed for the lysine probe (MiR/Parental = 1.78, p-value=0.0305, and MiR+Tram/Parental = 2.15, p-value=0.0098) (Figure 4E and 4F, Figure S11C and S11D). Therefore, in the BRAFi and MEK1/2i resistant cell lines, we observed a marked increase in overall probe reactivity, consistent with increased accessibility of one or more reactive cysteine and lysine residues within the respective resistance-associated proteins (Figure 4C-F) that was also observed at the 24-hour time point (Figure S6E). Because BiR cells overexpress BRAF, the elevated signal likely reflects greater total recovery of reactive protein. For MiR cells, the MEK2 Q60P mutation disrupts the inhibitory helix A interaction and stabilizes an active, open conformation, which may increase probe access to multiple reactive residues. Overall, these data demonstrate that ABPP detects composite reactivity shifts associated with inhibitor resistance—whether arising from protein overexpression or mutation-driven accessibility changes—without relying on single-residue assignment. Future studies will be necessary to determine whether these changes reflect altered accessibility at the same residues or increased labeling of previously inaccessible sites. Nonetheless, these findings provide a strong foundation for further investigation into how resistance-associated mutations affect kinase reactivity to chemo-proteomic probes and influence response to FDA-approved therapies.

**Figure 4.**
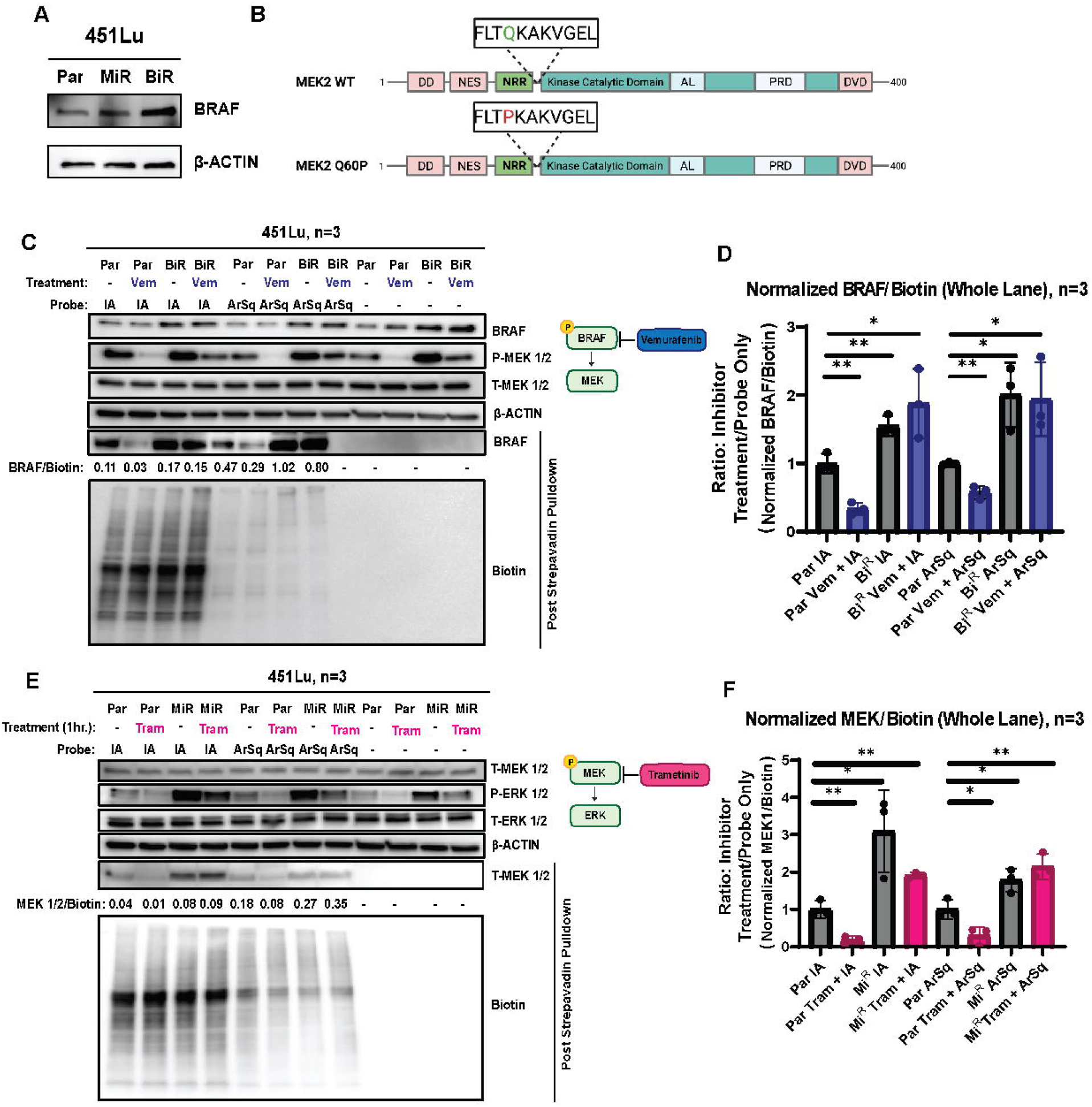
Resistance to MAPK pathway inhibitors induces distinct changes in cysteine and lysine accessibility. (A) Mechanisms of MAPK pathway inhibitor resistance in 451-Lu melanoma cells. Resistance to BRAF inhibitors is driven by overexpression of BRAF. Immunoblot detection of BRAF in 451-Lu parental (Par) cells and resistant derivatives, 451-Lu BRAFi^R^ (BiR) and 451-Lu MEKi^R^ (MiR). (B) Resistance to MEK1/2 inhibitors is conferred by a Q60P activating mutation in MEK2. (C) Immunoblot detection of BRAF, phosphorylated (P)-MEK1/2, total (T)-MEK1/2,, and β-actin, as well as BRAF and Biotin post streptavidin enrichment from the indicated cells treated with vehicle, 100 nM trametinib, or 1μM Vemurafenib for one hour. (D) Quantification: BRAF band intensity normalized to whole-lane Biotin and then to vehicle control. Results were compared using a pair-wise t-tests. One asterisk, P<0.05, Two asterisks, P<0.01, Three asterisks, P<0.001, Four asterisks, P<0.0001. n=3. (E) Immunoblot detection of T-MEK1/2, phosphorylated (P)-ERK1/2, total (T)-ERK1/2, and β-actin, as well as MEK1/2 and biotin after streptavidin pull-down from the indicated cells treated with vehicle, 100 nM trametinib, or 1μM Vemurafenib for one hour and labeled for one hour *in situ* with no probe, 25μM IA-alkyne, or 25μM ArSq-alkyne followed by Cu-catalyzed click chemistry with biotin azide. (F) Quantification: MEK1/2 band intensity normalized to whole-lane Biotin and then to vehicle control. Results were compared using pair-wise t-tests. One asterisk, P<0.05, Two asterisks, P<0.01, Three asterisks, P<0.001, Four asterisks, P<0.0001. n=3.

### Proteome-Wide Profiling Reveals Therapy-Induced Accessibility Changes Beyond Direct Drug Targets

Finally, we postulated that proteome-wide reactivity changes in response to kinase inhibition could reveal molecular features of therapeutic response, adaptation, and tumor cell vulnerabilities. To determine whether inhibitor treatment induces broader proteomic remodeling, we combined probe labeling with tandem mass tag (TMT)–based quantitative mass spectrometry. Proteins showing a > 1.5-fold increase or < 0.66-fold decrease in labeling with p < 0.05 were considered significantly changed. In A375 cells, trametinib altered cysteine reactivity in 23 proteins and vemurafenib in 86 proteins, while lysine reactivity changed in 74 and 50 proteins, respectively (Figure 5A). Although BRAF^V600E^ and MEK1/2 were not recovered—likely due to low abundance—these datasets included 270 kinases (Table S6), demonstrating broad kinome coverage [36]. Notably, the pseudokinase KSR2 exhibited one of the strongest decreases in cysteine labeling (2.6-fold reduction, p = 0.0415) following vemurafenib treatment (Figure 5A–C). Given that many kinases have relatively low abundance, we did not achieve significant coverage of many of these kinases suggesting that they may have been missed by traditional site directed ABPP. This highlights the power of using a protein level analysis to discover potential changes, before following up with additional studies on proteins of interest.

**Figure 5.**
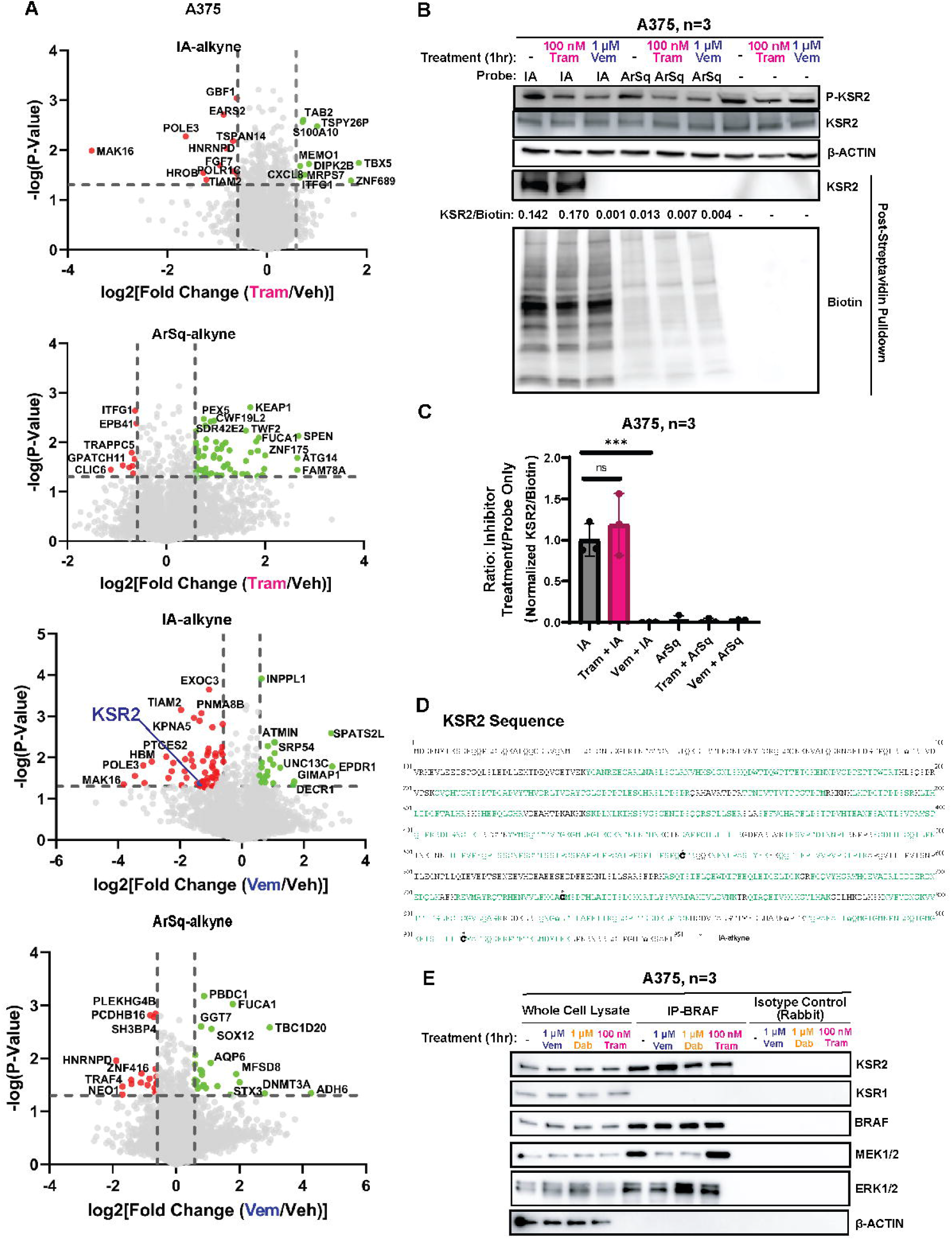
Therapy-induced changes in protein accessibility revealed by global chemoproteomic profiling. (A) Volcano plots display quantitative changes in peptide labeling post-streptavidin enrichment, as detected by mass spectrometry. Plots show –log10(p-value) versus log2(fold change), comparing 24 hour inhibitor-treated A375 cells to vehicle control. Top : Cysteine (IA-alkyne, 25 µM, one hour) accessibility changes following 100 nM trametinib treatment. Middle Top : Lysine (ArSq-alkyne, 25 µM, one hour) accessibility changes following 100 nM trametinib treatment. Middle Bottom : Cysteine (IA-alkyne, 25 µM, one hour) accessibility changes following 1 µM Vemurafenib treatment. Bottom : Lysine (ArSq-alkyne, 25 µM, one hour) accessibility changes following 1 µM Vemurafenib treatment. (B) Immunoblot detection of phosphorylated (P)-KSR2, total (T)-KSR2,and β-actin, as well as KSR2 and biotin after streptavidin pull-down from the indicated cells treated with vehicle, 100 nM trametinib, or 1μM Vemurafenib for one hour. in response to Trametinib and Vemurafenib treatment (C) Quantification: KSR2 band intensity normalized to whole-lane Biotin and then to vehicle control. Results were compared using pair-wise t-tests. One asterisk, P<0.05, Two asterisks, P<0.01, Three asterisks, P<0.001, Four asterisks, P<0.0001. n=3. (D) Sequence Map for KSR2 with annotated sites of labeling from IA-alkyne identified in overexpressed FlagKSR2. (E) Immunoblots of Co-IP of BRAF with MEK1/2, KSR2, KSR1, and ERK1/2, with β-actin for loading control of input.

Because KSR2 scaffolds MEK and ERK in MAPK signaling, we examined whether its decreased labeling reflected protein loss or altered accessibility. After one-hour inhibitor treatments, total KSR2 levels remained constant, but IA-alkyne labeling decreased significantly upon vemurafenib treatment (Figure 5B–C, Figure S12). In contrast, after a 24-hour treatment with vemurafenib, immunoblotting after a streptavidin pulldown showed a lack of cysteine accessibility on KSR2 (Figure S6G), but at this time point, immunoblotting in whole cell lysate showed lower KSR2 protein levels in vemurafenib treated conditions, which could account for the lack of probe labeling (Figure S6F). Additionally, due to the potential for other unintended consequences of a 24-hour treatment beyond the specific effect of vemurafenib treatment, we repeated the experiment at a one hour timepoint. We performed immunoblotting of whole-cell lysates to assess KSR2 expression and phosphorylation. Protein levels of KSR2 remained unchanged after one hour of inhibitor treatment, and phosphorylation levels decreased as expected following treatment with 1 µM vemurafenib or 100 nM trametinib (Figure 5B). To validate these findings, we performed streptavidin pulldown followed by immunoblotting and confirmed that IA-alkyne reactivity of KSR2 was reduced in A375 cells treated with vemurafenib (Figure 5B and 5C, Figure S12A and B). Site-level analysis of FLAG-KSR2 identified labeling at C552, C729, and C910 (Figure 5D). Reduced labeling at these sites could reflect local environment changes or altered complex assembly rather than direct binding effects. Additionally, across three replicates of these immunoblotting experiments, we detected no change in cysteine availability on KSR2 with trametinib treatment (Normalized MEK1/Biotin, compared to Veh: 1.19, p-value=0.418) (Figure 5C, Figure S12A and B). Collectively, these findings demonstrate that our chemoproteomic approach, combining activity-based probe labeling with TMT-based quantitative mass spectrometry, effectively reveals changes in residue accessibility following kinase inhibition. In particular, the reduction in KSR2 cysteine reactivity upon vemurafenib treatment underscores the potential of this strategy to uncover distinct molecular adaptations that drive therapeutic response and tumor cell vulnerabilities.

After confirming via immunoblotting that total KSR2 levels are unchanged after one hour of inhibitor treatment while IA-alkyne probe labeling of KSR2 decreases, we wanted to determine the role of protein-protein interactions in our observed changes in cysteine reactivity. Consistent with this, co-immunoprecipitation experiments showed that vemurafenib increased the association of KSR2 with BRAFV600E and MEK1/2, suggesting that modified protein–protein interactions may restrict probe access (Figure 5E). Additionally, vemurafenib treatment increased the KSR2 immunoprecipitated with BRAF^V600E^, indicating at least partially that protein-protein interactions could be responsible for our observed probe changes in BRAF^V600E^. The role of changing protein-protein interactions between KSR2 and BRAF^V600E^ as a result of vemurafenib treatment is furthered by the results of a streptavidin pulldown from A375 cells transiently overexpressing FLAG-KSR2 (Figure S13). After treating these cells for one hour with 1 μM vemurafenib and then probing *in situ* with IA-alkyne, similar to in Figure 5B, we observed an increase in FLAG-KSR pulldown compared to the control treatment. This led us to hypothesis that the overexpression of KSR2 resulted in levels that were outside of the normal stoichiometric range with its binding partners of BRAF and MEK. If our observations in endogenous KSR2 are to be made in line with our exogenous overexpressed KSR2, protein-protein interactions are a likely explanation for our observed changes in probe reactivity, though alternative explanations like changes in activity or conformational state cannot be definitively ruled out given the possibility that residues not covered in our site of labelling experiments could also contribute. Finally, STRING network analysis of proteins with reduced cysteine or lysine reactivity revealed enrichment in cell-cycle and division pathways (Figure S14), aligning with the known role of MAPK signaling in proliferation. Together, these proteome-level results extend ABPP’s utility beyond direct kinase targets, revealing composite residue-reactivity signatures that capture inhibitor-induced remodeling across signaling networks [29,30].

## Discussion

This study introduces a chemoproteomic framework for assessing inhibitor-associated changes in residue accessibility across key nodes of the MAPK signaling cascade. By combining amino acid–selective probes with short-term drug treatments and quantitative proteomics, we provide a practical approach for monitoring functional consequences of kinase inhibition in live cells without relying solely on phospho- or expression-based readouts. To establish a foundation for evaluating reactivity across a broad swathe of the proteome simultaneously, in response to targeted therapies, we first used cysteine- and lysine-reactive activity-based probes in combination with vemurafenib and trametinib, which target BRAF^V600E^ and MEK, respectively. We confirmed effective target engagement via MAPK signaling inhibition and verified that observed probe reactivity changes were not due to altered protein expression (Figure 2). Our initial in-gel fluorescence imaging demonstrated strong probe labeling across the proteome and revealed broad changes in residue accessibility, reinforcing the utility of this approach for capturing alterations to covalent probe reactivity (Figure S2). Immunoblotting of streptavidin-enriched samples revealed decreased cysteine and lysine accessibility in BRAF^V600E^ and MEK1/2 upon inhibitor treatment, consistent with inhibitor binding altering residue reactivity. Importantly, reversing the order of treatment—adding probes before inhibitors—abolished this effect, supporting the idea that conformational changes driven by inhibitor binding modulate accessibility. For MEK1, the reactive cysteine corresponds to C207 (Figure 2G and 2H), a site previously shown to influence copper binding and MEK1/2 activation [21,31]. Trametinib-induced decreases in lysine accessibility may reflect broader conformational changes, as no lysine residues are known to reside near its allosteric binding site [31]. Although we identified sites of lysine reactivity on MEK1 *in situ* with from ArSq-alkyne, we were unable to identify lysine adducts from the overexpression system or differences in either context when trametinib treatment was included, restricting our ability to provide mechanistic underpinning to our overall observation of decreased lysine accessibility via immunoblotting. For BRAF^V600E^, vemurafenib also reduced accessibility of cysteine and lysine residues in composite immunoblotting pulldown studies. These findings highlight how ABPP can detect conformational and binding-induced changes at reactive residues [32]. These findings expand on previous literature where ABPP has been utilized to demonstrate decreases in probe binding in response to covalent inhibitor engagement [8]. Our results demonstrate an application in live cells where we can use probe binding as an indicator for direct non-covalent inhibitor engagement. Therefore, ABPP can serve as an orthogonal method of determining targeted kinase inhibition without relying solely on the observation of downstream signaling changes. As probe design improves over time, a landscape for activity-based protein profiling for the detection of targeted changes in kinase activity, conformation states, and protein-protein interactions can improve on our collective understanding kinase functionality, particularly in systems where aberrant kinase activity drives negative disease outcomes.

While MEK1/2 and BRAF were not recovered in our TMT-based mass spectrometry dataset due to low abundance, we identified 234 proteins with significant changes in cysteine or lysine accessibility. Among these, KSR2 emerged as a compelling candidate due to its known role in MEK1/2 activation [37]. Vemurafenib decreased cysteine reactivity in endogenous KSR2, confirmed by both mass spectrometry and immunoblotting. This suggests a potential previously unrecognized interaction between vemurafenib and KSR2, possibly via conserved cysteines in the kinase-like domain, and raises new hypotheses about KSR complex formation in MAPK signaling [38-40]. These findings provide additional insight into existing literature that have sought to determine the role of MAPK pathway activation states with complex formation with KSR1 and KSR2 [41,42. Specifically, because BRAFV600E activation of MEK1/2 has been thought to have been facilitated by KSR1, our findings of immunoprecipitating KSR2 with BRAF but not KSR1 provides additional information on KSR complex assembly while in live cells in response to inhibitor treatments. Future work will be focused on more complete and quantitative changes to KSR2 reactivity with a variety of different probes to fully elucidate this mechanism.

Additionally, to assess the sensitivity of our approach to different inhibitor binding modes, we compared vemurafenib and dabrafenib—originally classified as type I and II BRAF^V600E^ inhibitors, respectively. Despite now both being categorized as type 1½ inhibitors, subtle differences in DFG-motif positioning remain as observed by previous crystal structures (Figure 3A). Using an aspartate/glutamate-reactive probe, we observed differential reactivity: vemurafenib decreased acidic residue accessibility, while dabrafenib increased it. These findings suggest probe-based profiling can resolve conformational shifts and support further use of acidic residue-reactive probes to study kinase dynamics [43]. Therefore, activity-based protein profiling techniques targeting key functional residues on kinases like lysines and aspartates can improve upon the current drawbacks that other approaches like immobilized inhibitor beads have had with failing to determine potential off-target binding of kinase inhibitors and determining kinase activation states. [44]

We extended our studies to drug-resistant models of BRAF^V600E^ melanoma using 451-Lu cells harboring either a Q60P mutation in MEK2 (MEKi-resistant) or BRAF amplification (BRAFi-resistant) [45,46]. As expected, both lines exhibited MAPK signaling persistence despite inhibitor treatment. Using our established probe workflow, we found that parental cells mirrored A375 patterns, with decreased probe labeling of MEK1 and BRAF^V600E^ following inhibitor treatment. In contrast, resistant cells showed increased probe accessibility for the corresponding resistant kinase. In BRAFi-resistant cells, this likely reflects BRAF overexpression, leading to greater pull-down. However, in MEKi-resistant cells, the Q60P mutation in MEK2 may drive conformational changes that increase reactive residue sidechain accessibility. Importantly, increased activity of MEK1 and BRAF could also result in the perceived increase in the targets post pull-down, through changes in substrate binding, active conformation, or interactions with other proteins. Though, given our observations on the sites of differential probing on the kinases in different activation states, it is interesting to note that without a different conformation due to the Q60P mutation, we would expect similar decreases in probe binding as in the 451-Lu Parental cell line, which is consistent with the observed changes in the A375 cell line. This suggests mutationally induced structural changes can create new reactive residues, offering opportunities for developing next-generation inhibitors that exploit these vulnerabilities. Overall, our study demonstrates that activity-based proteome profiling is a powerful approach to monitor kinase conformational states, assess treatment response, and uncover adaptive vulnerabilities in drug-resistant cancer.

## Conclusion

In this study, we employed amino acid–specific chemical probes targeting cysteine, lysine, and acidic residues (aspartate and glutamate) to investigate accessibility changes in BRAF^V600E^ melanoma cells in response to MAPK pathway inhibition. We identified consistent decreases in cysteine and lysine accessibility following treatment with vemurafenib and trametinib, as well as differential labeling of BRAF aspartate/glutamate residues when comparing vemurafenib to dabrafenib, reflecting potential conformational distinctions between the two inhibitor/protein complexes. In resistant melanoma cell lines, we observed marked increases in residue accessibility on MEK and BRAF, consistent with inhibitor-induced conformational remodeling associated with acquired resistance. Finally, mass spectrometry–based profiling uncovered additional changes in reactivity in proteins beyond the known drug targets, including KSR2, highlighting broader proteomic adaptations to MAPK pathway inhibition.

