## Supplementary Information for "MAPK Pathway Inhibition Reshapes Kinase Chemical Probe Reactivity Reflecting Cellular Activation States"

### Supplemental Figures

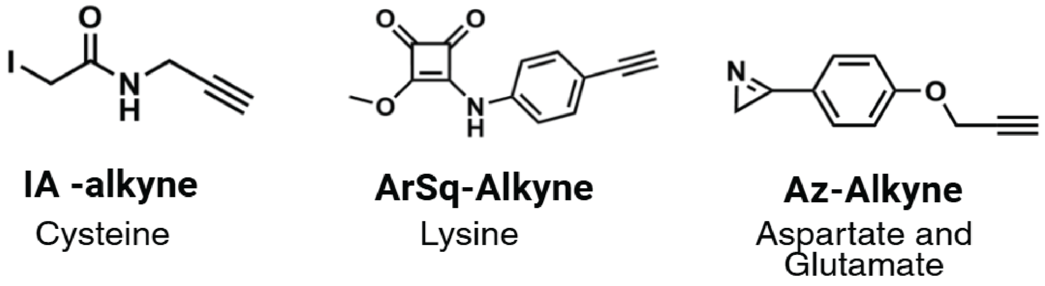


**Supplemental Figure 1.** Chemical structures of amino acid–reactive probes used in this study. Left: IA (iodoacetamide)-alkyne; Middle: ArSq-alkyne; Right: Az-alkyne.

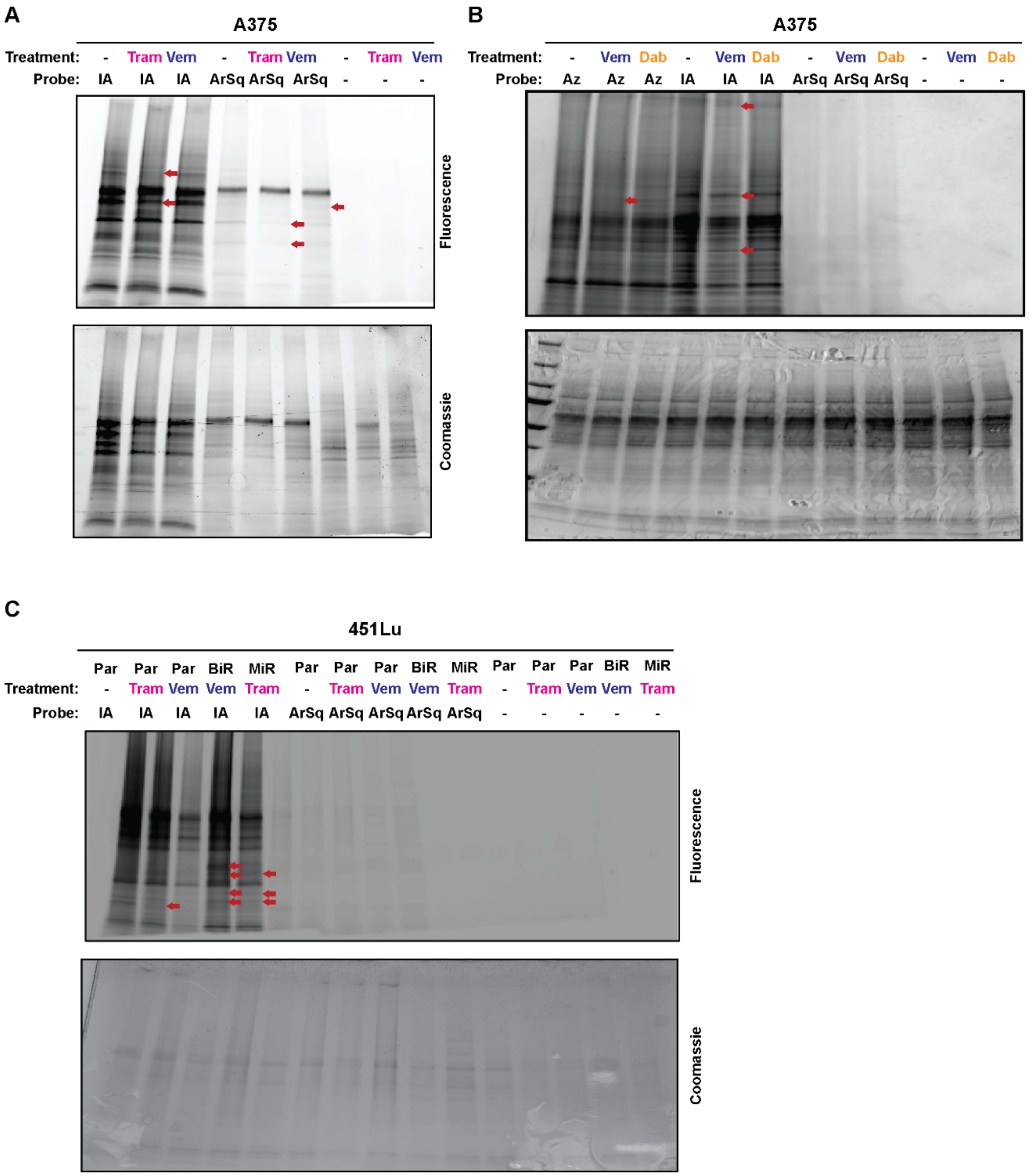


**Supplemental Figure 2.** In-gel fluorescence and Coomassie stained gels from (A) Figure 2D A375 cells treated with vehicle, 100 nM trametinib, or 1μM vemurafenib, for 24 hours and labeled with no probe, IA-alkyne, or ArSq-alkyne followed by Cu-catalyzed click chemistry with TAMRA-biotin azide. (B) Figure 3B A375 cells treated with vehicle, 1μM vemurafenib, or 1μM dabrafenib, for 24 hours and labeled with no probe, AZ-alkyne, IA-alkyne, or ArSq-alkyne followed by Cu-catalyzed click chemistry with TAMRA-biotin azide, and (C) Figure 4C indicated cells treated with vehicle, 100 nM Trametinib, or 1μM Vemurafenib for 24 hours and labeled with no probe, IA-alkyne, or ArSq-alkyne followed by Cu-catalyzed click chemistry with TAMRA-biotin azide. Changes in band intensity in inhibitor conditions are highlighted using red arrows.

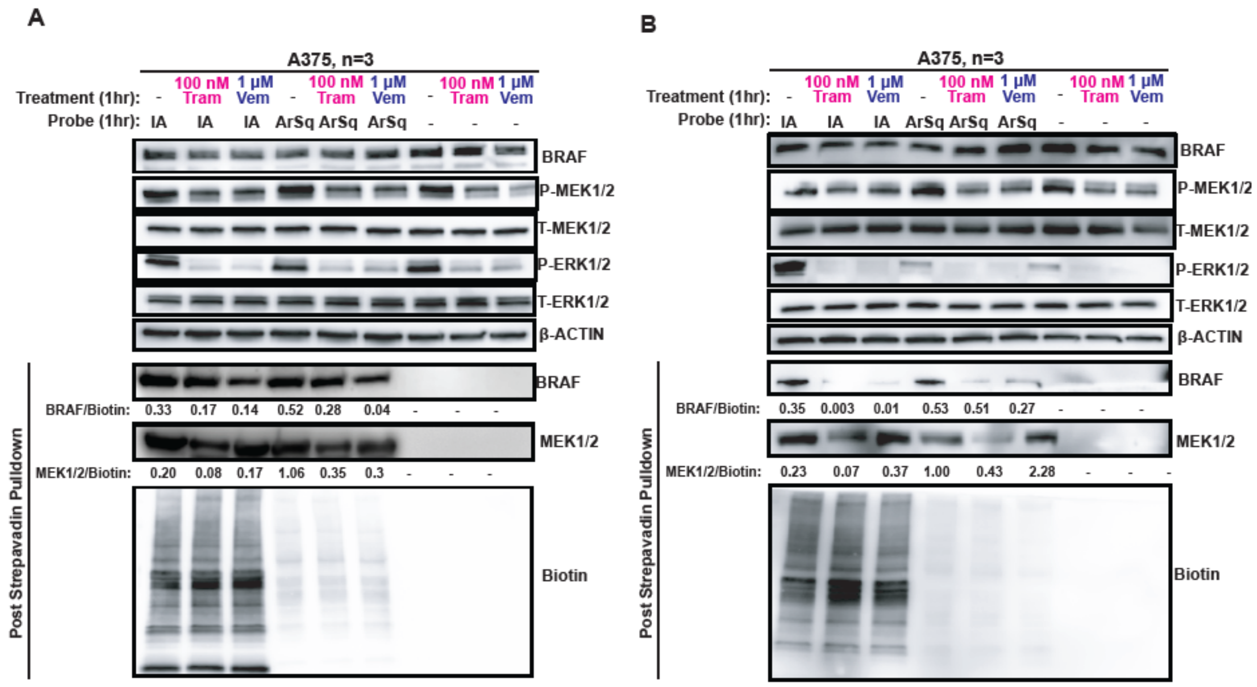


**Supplemental Figure 3. Replicates of Streptavidin Pulldown Immunoblots from Figure 2C.** (A) and (B) Immunoblot detection of BRAF, phosphorylated (P)-MEK1/2, total (T)-MEK1/2, P-ERK1/2, T-ERK1/2, and b-actin from A375 cells treated for 1 hour with vehicle, 100 nM trametinib, or 1μM vemurafenib, and labeled in-situ for 1 hour with no probe, IA-alkyne, or ArSq-alkyne. Additional immunoblots of BRAF, MEK 1/2 and Biotin after streptavidin-enrichment, including densitometry quantification normalizing BRAF or MEK1/2 post streptavidin enrichment to Whole Lane Biotin, corresponding to the results of Figure 2C.

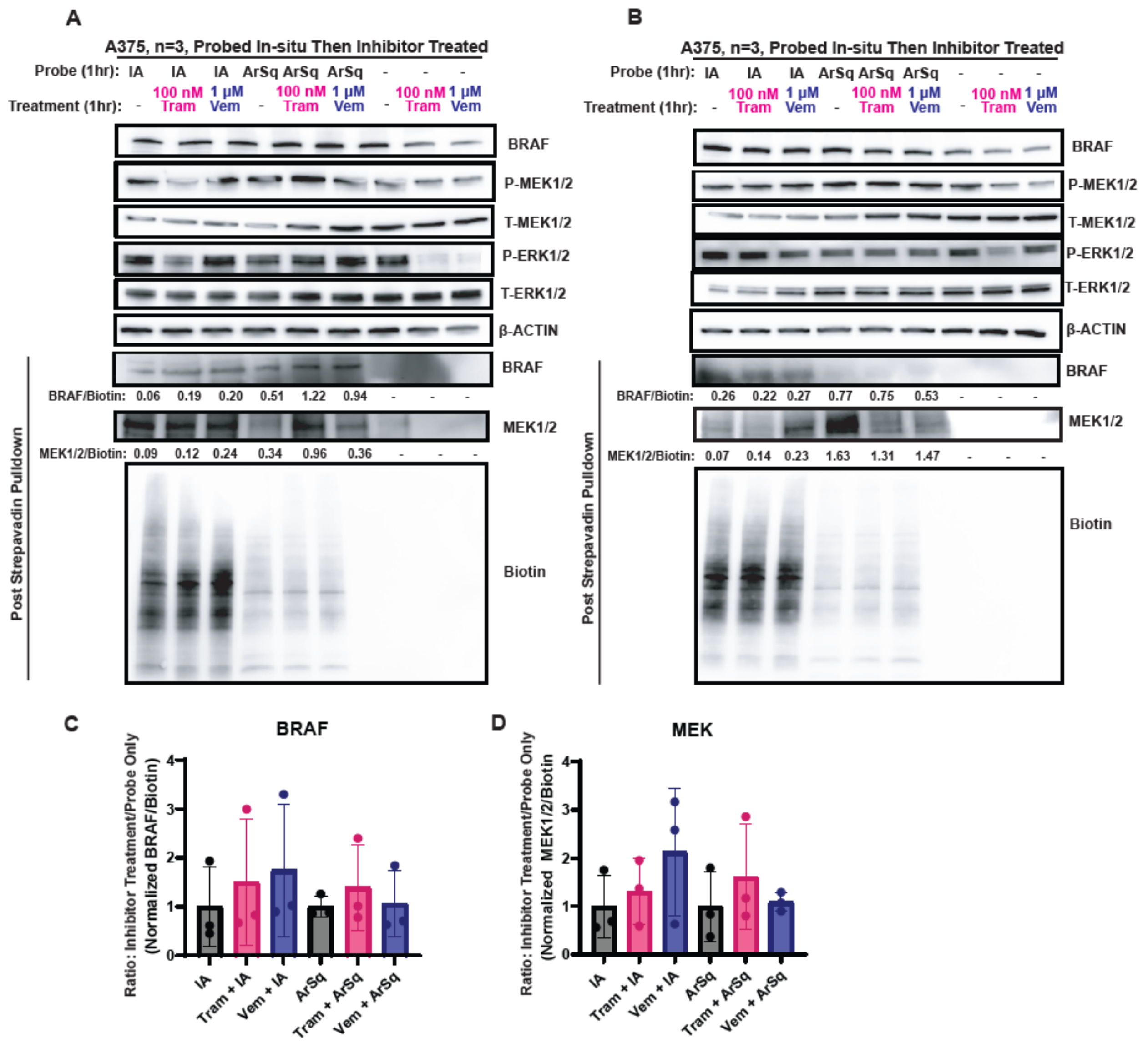

**Supplemental Figure 4. Replicates of Streptavidin Pulldown Immunoblots from Figure 2D.** A) and (B) Immunoblot detection of BRAF, phosphorylated (P)-MEK1/2, total (T)-MEK1/2, P-ERK1/2, T-ERK1/2, and b-actin from A375 cells labeled in-situ for 1 hour with no probe, IA-alkyne, or ArSq-alkyne and then treated for 1 hour with vehicle, 100 nM trametinib, or 1µM vemurafenib. Additional immunoblots of BRAF, MEK 1/2 and Biotin after streptavidin-enrichment, including densitometry quantification normalizing BRAF or MEK1/2 post streptavidin enrichment to Whole Lane Biotin, corresponding to the results of Figure 2D. (C) Densitometry Quantification: BRAF band intensity normalized to Biotin and then to vehicle control. (D) Densitometry Quantification: MEK1/2 band intensity normalized to Biotin and then to vehicle control.

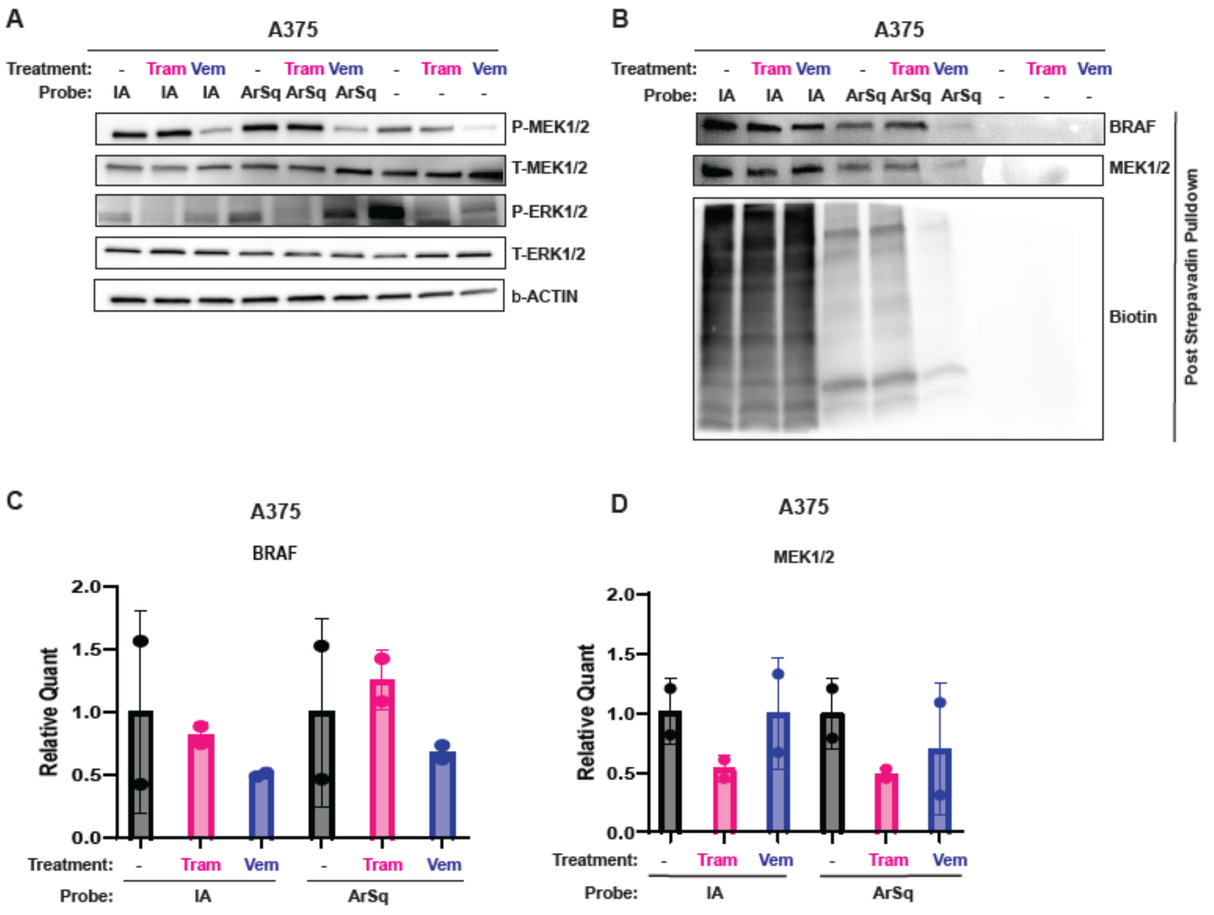


**Supplemental Figure 5. In-lysate probe labeling reveals similar changes in cysteine and lysine reactivity of MEK1/2 and BRAF following inhibitor treatment.** (A) Immunoblot detection of phosphorylated (P)-MEK1/2, total (T)-MEK1/2, P-ERK1/2, T-ERK1/2, and b-actin from A375 cells treated with vehicle, 100 nM trametinib, or 1μM temurafenib, for 24 hours and in-lysate labeling with no probe, IA-alkyne, or ArSq-alkyne. (B) Immunoblot detection of BRAF, MEK1/2, or biotin after streptavidin pull-down from A375 cells treated with vehicle, 100 nM Trametinib, or 1μM Vemurafenib, for 24 hours and in-lysate labeling with no probe, IA-alkyne, or ArSq-alkyne by Cu-catalyzed click chemistry with biotin azide. (C) Densitometry Quantification: BRAF band intensity normalized to Biotin and then to vehicle control. (D) Densitometry Quantification: MEK1/2 band intensity normalized to Biotin and then to vehicle control.

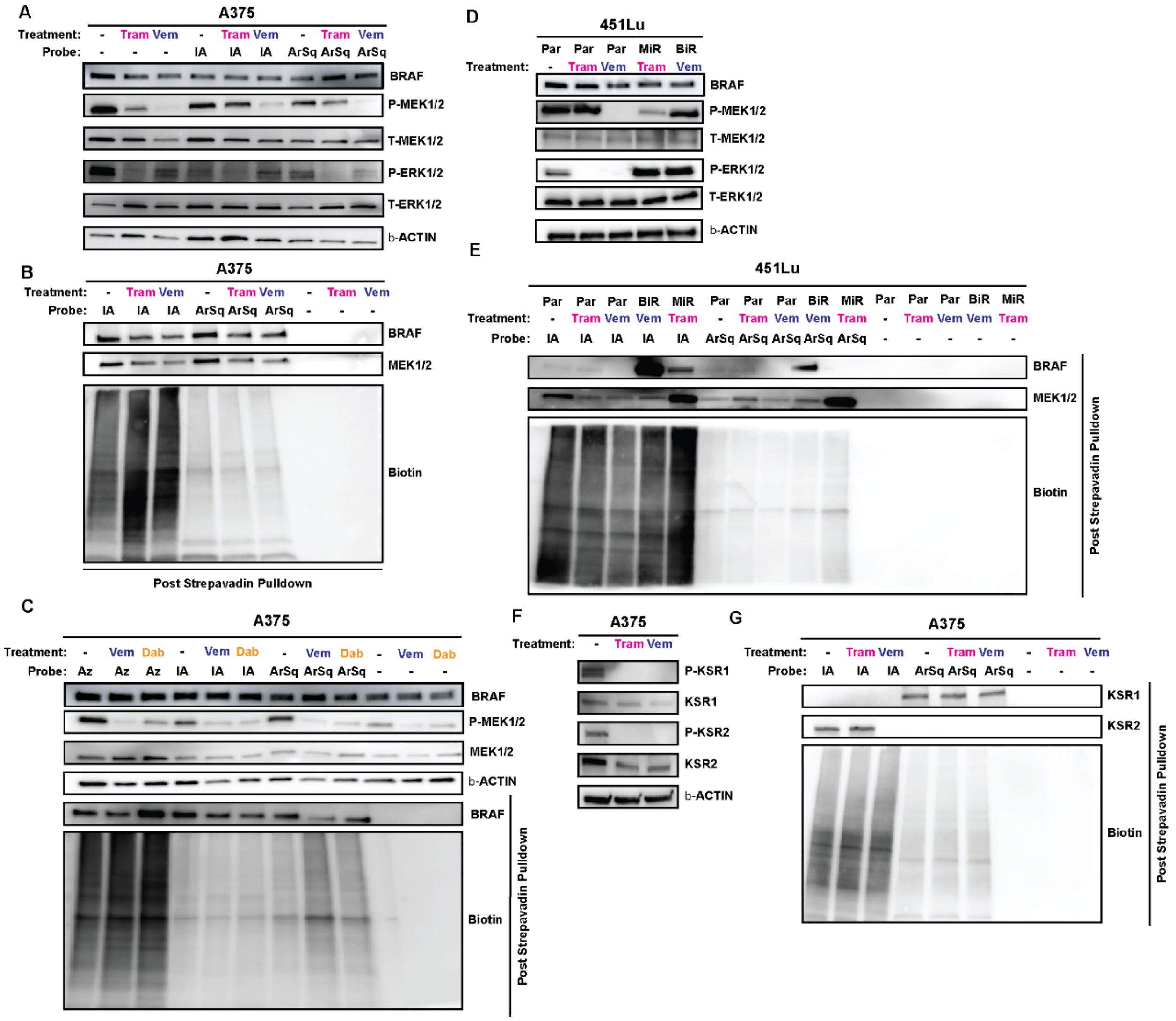


**Supplemental Figure 6. Assessment of accessibility changes in cysteine, lysine, and/or carboxylic acid residues of MEK, BRAF, and KSR2 after 24-hour treatment with targeted inhibitors for MEK and BRAF.** (A) Immunoblot detection of phosphorylated (P)-MEK1/2, total (T)-MEK1/2, P-ERK1/2, T-ERK1/2, and b-actin from A375 cells treated for 24 hours with vehicle, 100 nM trametinib, or 1μM vemurafenib, and labeled in-situ for 1 hour with no probe, IA-alkyne, or ArSq-alkyne, corresponding to the results from Figure 2D. (B) Immunoblots of BRAF, MEK 1/2 and Biotin after streptavidin-enrichment according to the same treatment scheme as in (A). (C) Immunoblot detection of phosphorylated (P)-MEK1/2, total (T)-MEK1/2, and b-actin from A375 cells treated with vehicle, 1μM vemurafenib, or 1μM dabrafenib for 24 hours and labeled for 1 hour with no probe, 10μM AZ-alkyne, 25μM IA-alkyne, or 25μM ArSq-alkyne. Immunoblot detection of BRAF or biotin after streptavidin pull-down from A375 cells treated with vehicle, 1μM vemurafenib, or 1μM dabrafenib, for 24 hours and labeled with no probe, AZ-alkyne, IA-alkyne, or ArSq-alkyne followed by Cu-catalyzed click chemistry with biotin azide. (D) Immunoblot detection of MEK 1/2 and ERK 1/2 phosphorylation in 451-Lu Parental, MEK-Inhibitor Resistant, and BRAF-Inhibitor Resistant cells after 24 hours of treatment with vehicle, 100nM

Trametinib, or 1µM vemurafenib. (E) Immunoblotting of BRAF, MEK1/2, and Biotin after streptavidin enrichment in 451-Lu Parental, MEK-Inhibitor Resistant, and BRAF-Inhibitor Resistant cells treated for 24 hours with vehicle, 100nM Trametinib, or 1µM vemurafenib, and then probed in-situ for one hour with vehicle, 25µM IA-alkyne, or 25µM ArSq-alkyne. (F) Immunoblot detection of P-KSR1, KSR1, P-KSR2, KSR2, and Beta-Actin in A375 cells treated for 24 hours with vehicle, 100nM Trametinib, or 1µM vemurafenib. (G) Immunoblot detection of KSR1, KSR2, and Biotin after streptavidin enrichment in A375 cells treated for 24 hours with vehicle, 100nM Trametinib, or 1µM vemurafenib, and then probed in-situ for one hour with vehicle, 25µM IA-alkyne, or 25µM ArSq-alkyne.

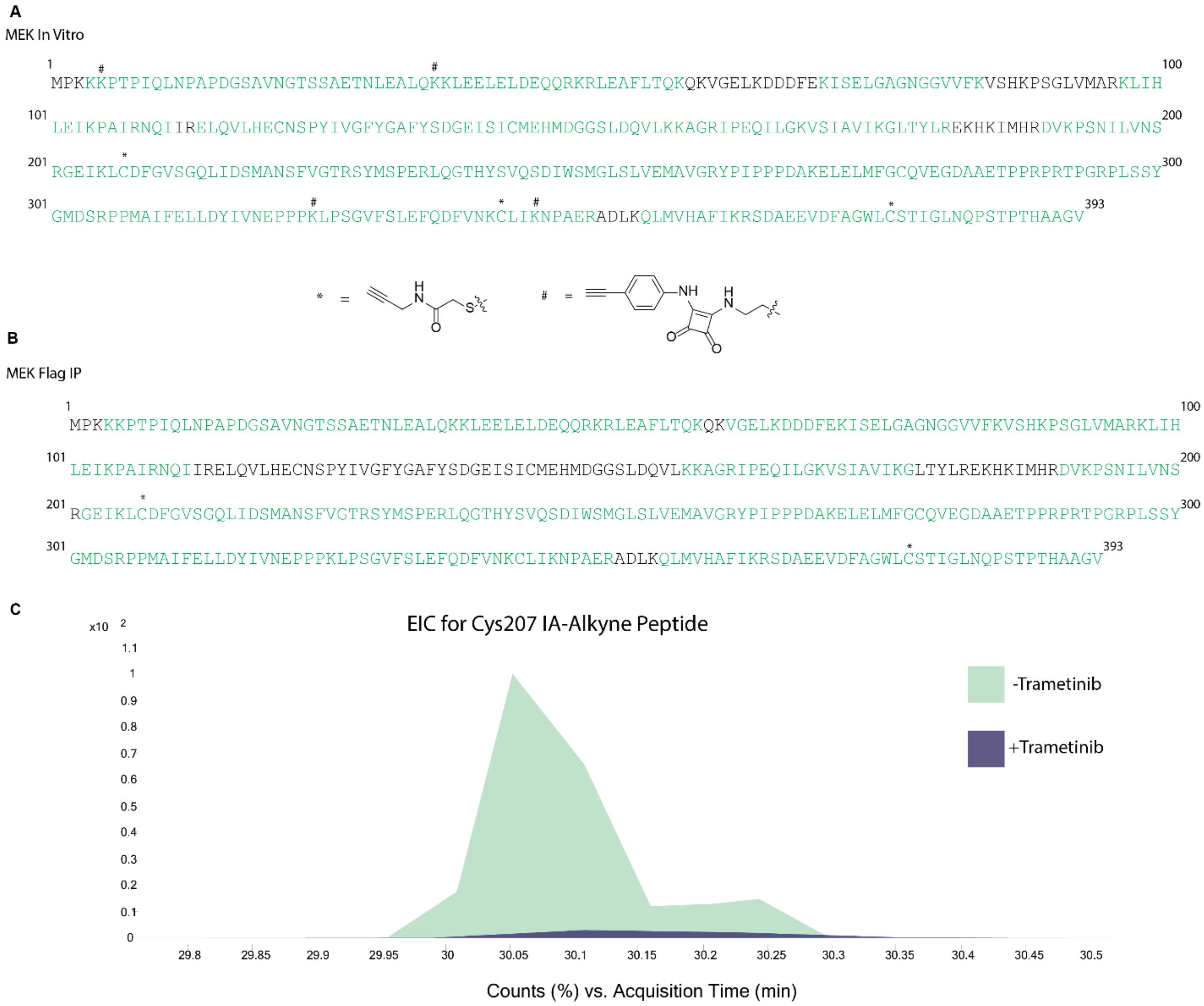


**Supplemental Figure 7. Sites of Labelling Observed on MEK1**. (A) Coverage Map of recombinant MEK1 with indicated sites of labelling (Green denotes all mapped peptides) (B) Coverage Map of FLAG-MEK1 immunoprecipitated from A375 cells with indicated sites of labelling (Green denotes all mapped peptides). (C) Extracted ion chromatogram for Cys207 IA-Alkyne peptide from recombinant MEK1 in the presence or absence of Trametinib.

BRAF In Vitro

416 LQKSPGPQRERKSSSSSEDRNRMKTLGRRDSSDDWEIPDGQITVGQRIGSGSFGTVYKGKWHGDVAVKMLNVTAPTPQQLQAFKNEVGVLRKTRHVNILL 515

516 FMGYSTKPQLAIVTQWCEGSSLYHHLHIIETKFEMIKLIDIARQTAQGMDYLHAKSIIHRDLKSNNIFLHEDLTVKIG**DFG**LATVKSRWSGSHQFEQLSG 615

616 SILWMAPEVIRMQDKNPYSFQSDVYAFGIVLYELMTGQLPYSNINNRDQIIFMVGRGYLSPDLSKVRSNCPKAMKRLMAECLKKKRDERPLFPQILASIE 715

716 LLARSLPKIHRSASEPSLNRAGFQTEDFSLYACASPKTPIQAGGYGAFPVH 766

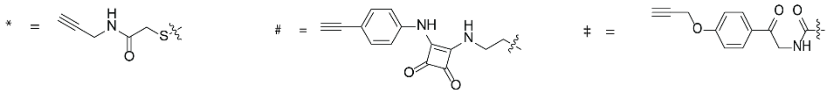


**Supplemental Figure 8. Sites of Labelling Observed on BRAF V600E (416-766).** (A) Coverage Map of recombinant BRAF V600E (416-766) with indicated sites of labelling (Green denotes all mapped peptides)

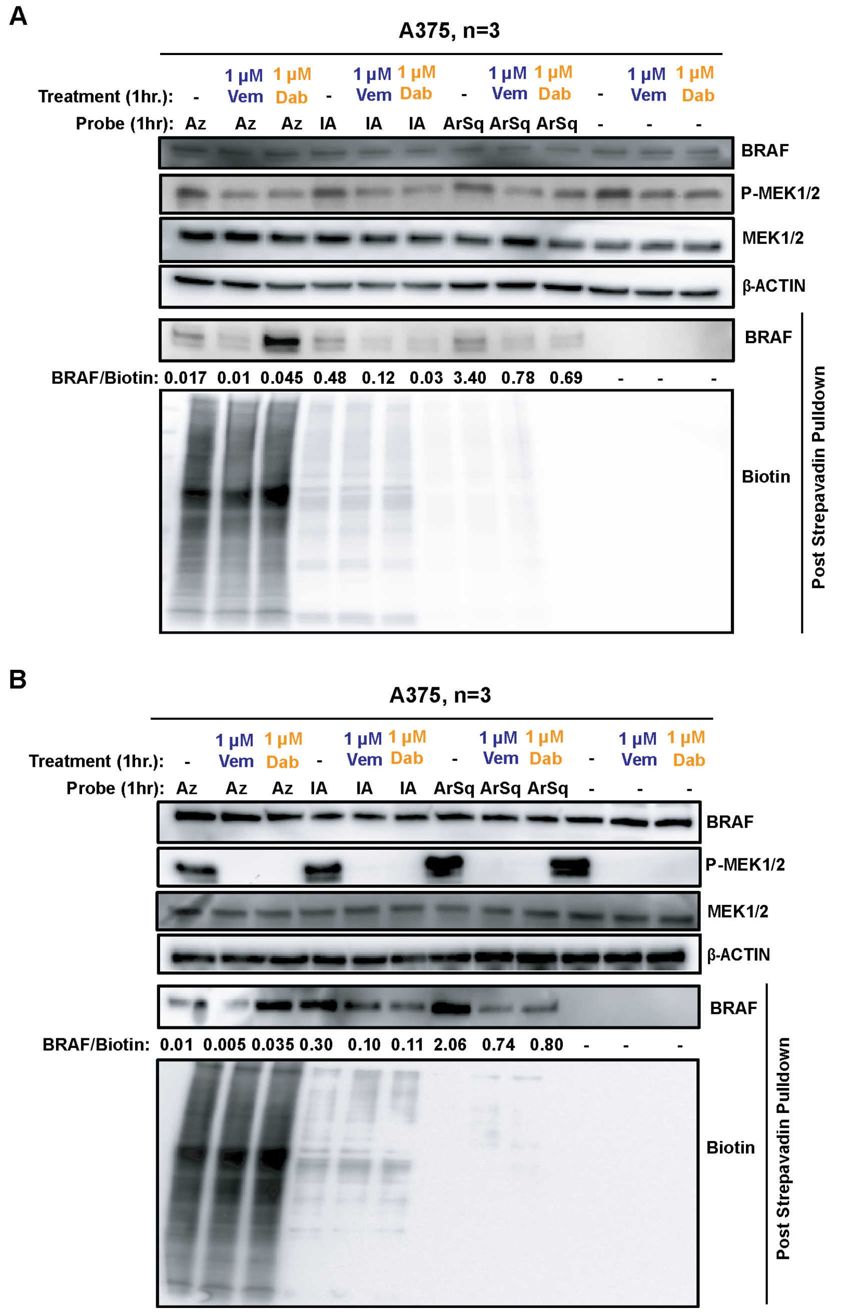


**Supplemental Figure 9. Replicates of Streptavidin Pulldown Immunoblots from Figure 3B.** (A) and (B) Immunoblot detection of BRAF, phosphorylated (P)-MEK1/2, total (T)-MEK1/2, and b-actin from A375 cells treated for 1 hour with vehicle, 1μM dabrafenib, or 1μM vemurafenib, and labeled in-situ for 1 hour with no probe, IA-alkyne, or ArSq-alkyne. Additional immunoblots of BRAF and Biotin after streptavidin-enrichment, including densitometry quantification normalizing post streptavidin enrichment BRAF to Whole Lane Biotin, corresponding to the results of Figure 3B.

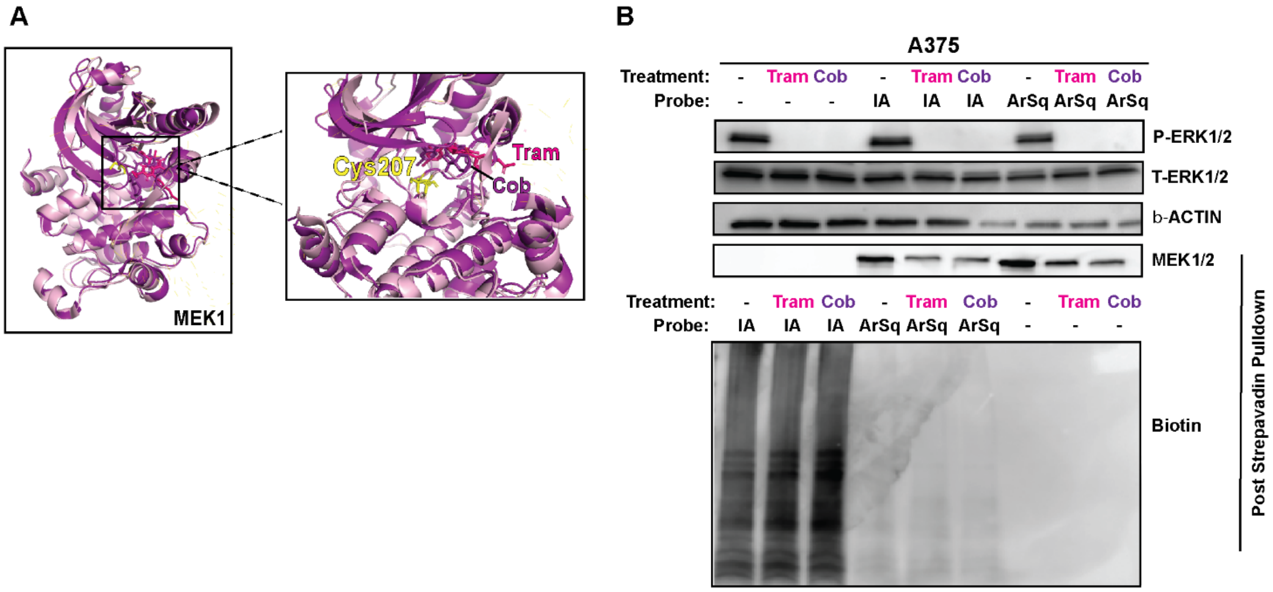


**Supplemental Figure 10. Comparison of cysteine and lysine ligandability in MEK1/2 following treatment with cobimetinib or trametinib.,** (A) Left: Crystal structure of cobimetinib bound to MEK1 overlayed with crystal structure of trametinib bound MEK1; Right: Zoom in to Trametinib (Pink) and Cobimetinib (Purple) binding site with C207 highlighted in yellow. (B) Immunoblot detection of phosphorylated (P)-ERK1/2, total (T)-ERK1/2, and β-actin from A375 cells treated with vehicle, 100 nM trametinib, or 100nM cobimetinib for 24 hours and in-lysate labeling with no probe, IA-alkyne, or ArSq-alkyne. Immunoblot detection of MEK1/2 or biotin after streptavidin pull-down from A375 cells treated with vehicle, 100 nM Trametinib, or 100nM cobimetinib or 24 hours and labeled with no probe, IA-alkyne, or ArSq-alkyne followed by Cu-catalyzed click chemistry with biotin azide.

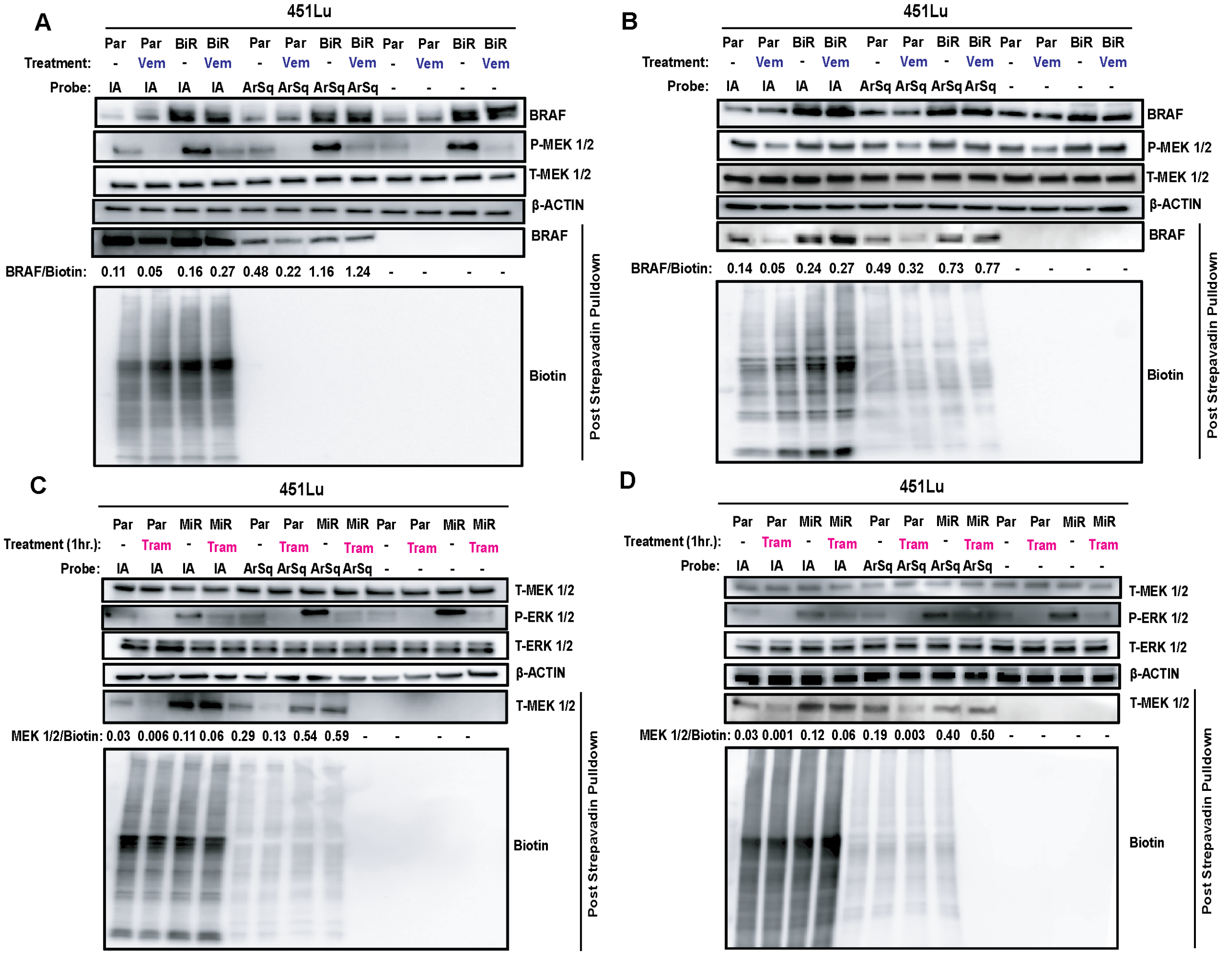


**Supplemental Figure 11. Replicates of Streptavidin Pulldown Immunoblots from Figure 4C and 4E.** (A) and (B) Immunoblot detection of BRAF, phosphorylated (P)-MEK1/2, total (T)-MEK1/2, and b-actin from 451-Lu Parental or BRAF-Inhibitor Resistant cells treated for 1 hour with vehicle or 1μM vemurafenib, and labeled in-situ for 1 hour with no probe, IA-alkyne, or ArSq-alkyne. Additional immunoblots of BRAF and Biotin after streptavidin-enrichment, including densitometry quantification normalizing post streptavidin enrichment BRAF to Whole Lane Biotin, corresponding to the results of Figure 4C. (C) and (D) Immunoblot detection of total (T)-MEK1/2, P-ERK1/2, T-ERK1/2, and b-actin from 451-Lu Parental or MEK-Inhibitor Resistant cells treated for 1 hour with vehicle or 100nM Trametinib, and labeled in-situ for 1 hour with no probe, IA-alkyne, or ArSq-alkyne. Additional immunoblots of MEK1/2 and Biotin after streptavidin-enrichment, including densitometry quantification normalizing post streptavidin enrichment MEK1/2 to Whole Lane Biotin, corresponding to the results of Figure 4E.

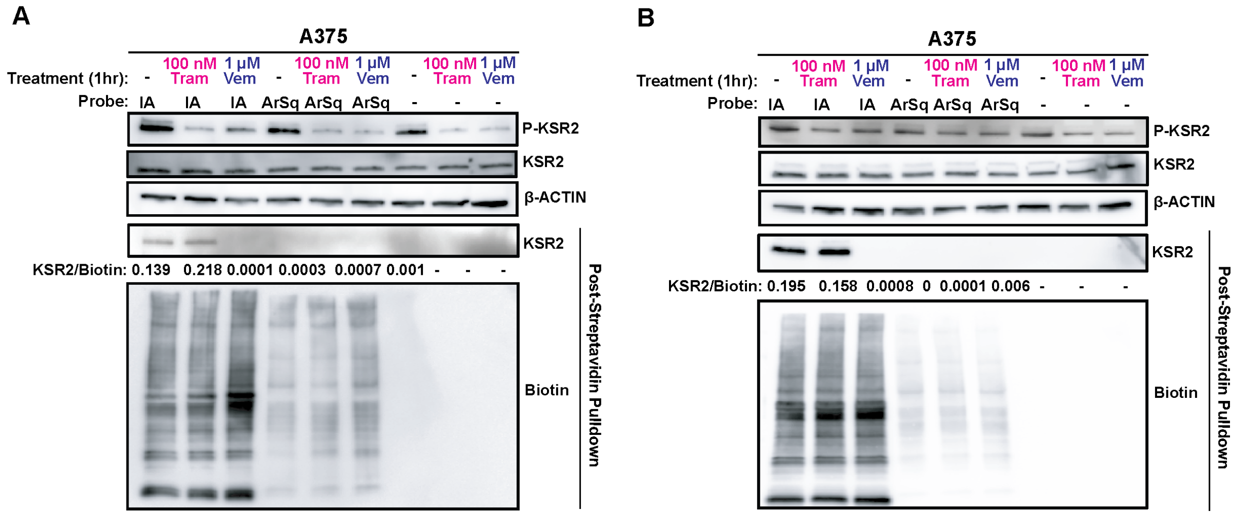


**Supplemental Figure 12. Replicates of Streptavidin Pulldown Immunoblots from Figure 5B.** (A) and (B) Immunoblot detection of phosphorylated (P)-KSR2, total (T)-KSR2, and b-actin from A375 cells treated for 1 hour with vehicle, 100 nM trametinib, or 1µM vemurafenib, and labeled in-situ for 1 hour with no probe, IA-alkyne, or ArSq-alkyne. Post-streptavidin enrichment immunoblots for T-KSR2 and Biotin were used for densitometry quantification of normalized KSR2 to Whole Lane Biotin, corresponding to the results of Figure 5B.

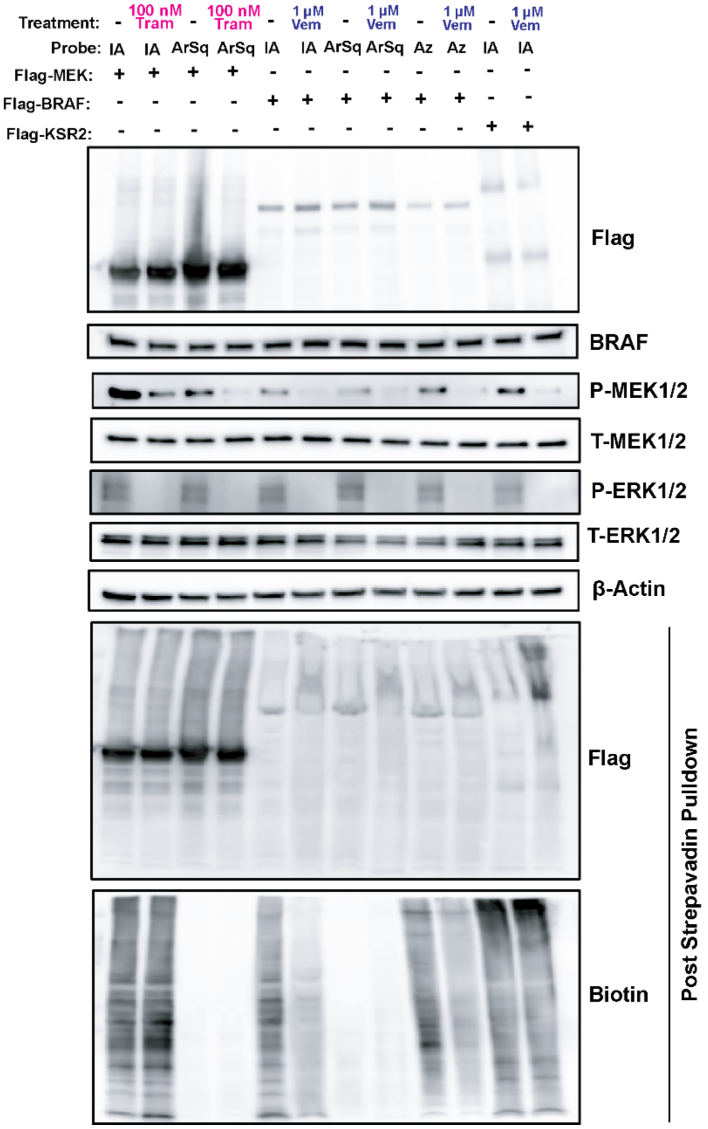


**Supplemental Figure 13. Streptavidin Enrichment from A375 cells exogenously expressing flag-tagged MEK1, BRAFV600E, and KSR2.** Immunoblot detection of Flag-tagged MEK1/2, BRAF, KSR2, phosphorylated (P)-MEK1/2, Total (T)-MEK1/2, phosphorylated (P)-ERK1/2, total (T)-ERK1/2, and b-actin from A375 cells expressing one of either flag-tagged MEK1, BRAFV600E, or KSR2. Flag-MEK1 A375 cells were treated for one hour with either vehicle or 100nM Trametinib and then probed in-situ for one hour with either 25µM IA-alkyne or 25µM ArSq-alkyne. Flag-BRAFV600E A375 cells were treated for one hour with either vehicle or 1µM vemurafenib and then probed in-situ for one hour with either 25µM IA-alkyne, 25µM ArSq-Alkyne, or 10µM Az-alkyne. Flag-KSR2 A375 cells were treated with vehicle or 1µM vemurafenib and then probed in-situ with 25µM IA-alkyne. Additional immunoblot detection of Flag-tagged MEK1, BRAFV600E, and KSR2 with Biotin post streptavidin pulldown

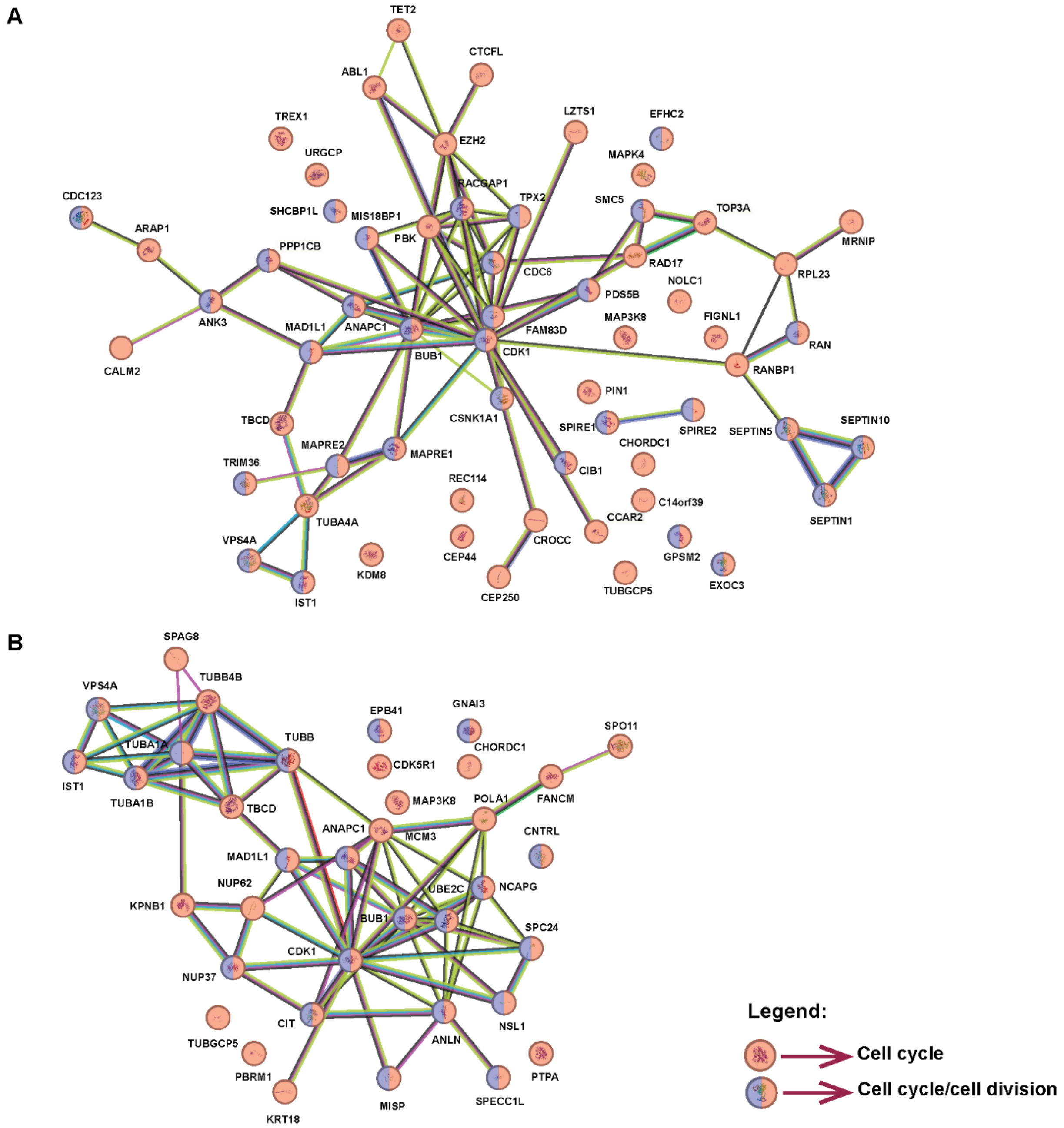


**Supplemental Figure 14. STRING analysis of proteins with reduced probe labeling following MAPK inhibitor treatment.** (A) Vemurafenib-treated A375 cells labeled with IA-alkyne and (B) trametinib-treated A375 cells labeled with ArSq-alkyne were analyzed by STRING for functional enrichment. Proteins with significantly decreased labeling were enriched for biological processes including cell cycle regulation, mitotic spindle organization, and cellular division.

### Materials and Methods

#### Cell Lines and Culture

293T/17 (ATCC, catalog #CRL-11268) and A375 (ATCC, catalog #CRL-1619) cells were purchased from the indicated companies and maintained in RPMI 1640 w/ Glutamax (Gibco, catalog #72400047) supplemented with 10% v/v fetal bovine serum (FBS, Cytvia, catalog #SH30071.03) and 1% penicillin-streptomycin (P/S, Gibco, catalog #15140122). 451Lu parental cells and resistant derivatives, 451-Lu BRAFi$^{R}$ and 451-Lu MEKi$^{R}$ [34,35] were a kind gift from Jessie Villanueva (Wistar Institute) and maintained in RPMI 1640 w/ Glutamax DMEM supplemented with 10% FBS with 1 µM vemurafenib or 1 µM trametinib (summarized in Table S1). Cell lines were not authenticated. Mycoalert testing was done to test for mycoplasma contamination of all cell lines.

#### Generation of A375 Cell Lines Exogenously Expressing Flag-Tagged MEK1, BRAFV600E, or KSR2

pcDNA3.1-FLAG-BRAFV600E and was pcDNA3.1-FLAG-KSR2 were created by Gateway recombination from pDONR223 BRAFV600E (Addgene plasmid #81700) into pcDNA3.1-3xFLAG-CCDB. pcDNA5-FLAG-MEK1 was created by Gibson assembly from pEGFP-C1-MEK1 with primers designed to include an N-terminal FLAG-tag. Briefly, a pcDNA5-FLAG vector fragment and a MEK1 fragment were amplified by PCR. DpnI at 37 ºC for 1 hour was used to digest template vectors. PCR products were separated by agarose gel electrophoresis and bands of the correct size were extracted (Qiagen QIAquick Gel Extraction Kit). Gibson assembly was performed in a 2:1 ratio of vector:insert using the HiFi DNA Assembly Master Mix, incubated for 50 ºC for 1 hour. A375 cells transiently expressing pcDNA3.1-FLAG-BRAFV600E or pcDNA5-FLAG-MEK1 were transfected using established protocols.

**Table S1: MAPK Pathway Inhibitors**

| Name | Source (Cat#) | Stock Concentration |
|---|---|---|
| Vemurafenib | ChemieTek (CT-P4032) | 1 mM |
| Dabrafenib | SelleckChem (S2807) | 1 mM |
| Trametinib | SelleckChem (S2673) | 100 µM |
| Cobimetinib | SelleckChem (S8041) | 100 µM |

#### Sample Collection and Preparation

One million of the indicated cells were plated in 10 cm dish (Genesee Scientific, catalog#25-202) and treated with either vehicle control (DMSO) or a MAPK inhibitor (summarized in Table S1) for either 1 or 24 hours. Treated cells were then washed with cold 1X PBS (phosphate buffered saline) and then collected via scraping in 1 mL of cold 1X PBS and transferred to 1.7 mL microcentrifuge tubes (Genesee Scientific, catalog#24-282). The cells were centrifuged at 2000 rpm for 3 minutes, and then resuspended in 500 µL of cold 1X PBS. Each resuspended cell pellet was then sonicated (Fisher Scientific, catalog# FB120110) for 2 rounds of 10 pulses for 1 second at 25% power with 10 minutes in between sonication rounds. The sonicated resuspended cell pellets were first centrifuged for 12,000 rpm for 5 minutes at $4^oC$ and then transferred to ultracentrifuge tubes (Labcon, catalog#3016-870-000)to be centrifuged at 45,000 rpm at $4^oC$ for 30 minutes in Sorvall MX120 Plus micro-ultracentrifuge to separate soluble and insoluble protein fractions. The soluble protein samples were transferred back to 1.7 mL microcentrifuge tubes. The protein concentration was determined by Peirce BCA Protein Assay Kit (Thermo Scientific, catalog#23227) using BSA as a standard. Protein lysates were processed for downstream click chemistry enrichment and subsequently analyzed via in-gel fluorescence, mass spectrometry, or immunoblotting.

Purified MEK1 (cat#:PR5043A) and BRAFV600E (cat#:PR7304A) Recombinant Human Protein were purchased From Thermo Scientific.

**Click Chemistry Probes**

Iodoacetamide-alkyne (cysteine) was purchased from Thermo-Scientific (catalog #I10189). ArSq-alkyne (lysine) and Az-alkyne (Aspartate and Glutamate) were synthesized according to the published procedures in Ivancova et. al and Ma *et* al. respectively [36,37].

The probes were first reconstituted in DMSO to make 100 mM aliquots. Each probe was then added directly to the cell culture media at a concentration of 25 µM, and then left to incubate at $37^oC$ for one hour.

**Click Chemistry Reaction**

The general click reaction mix per 25 µL reaction volume was: 1.5 µL of 1.7 mM Tris-benzyltriazolylmethyl-amine (TBTA, Cayman Chemical Company, catalog #18816, in DMSO), 0.5 µL of 50mM $CuSO_4$ (Sigma-Aldrich, catalog #C8027-500G, in Milli-Q $H_2O$), 0.5 µL of 13 mg/mL Tris(2-carboxyethyl)phosphine hydrochloride (TCEP, Sigma-Aldrich, catalog #C4706-2G, in $H_2O$), and 0.5 uL of 100 mM Biotin Azide (Specifications below for specific applications). The master mix reagents are added in the order listed above to each respective sample and incubated at room temperature for one hour.

After normalizing protein concentration, the sample was split into 25 µg, 250 µg and 1mg fractions. The 25 µg fraction was click labelled with a TAMRA-Biotin Azide (Vector Laboratories, catalog #CCT-1048, in DMSO) for in-gel fluorescence. The remaining fractions were click labelled with Azide-PEG3-Biotin conjugate (Sigma-Aldrich, catalog #762024, in DMSO) for Immunoblotting and mass spectrometry post streptavidin pulldown.

**In-Gel Fluorescence**

After one hour incubation with click-reaction mix, 5x Sample Buffer (0.25 M Tris pH 6.8 Tris, 10% SDS, 50% Glycerol, 25% β-Mercaptoethanol, 0.5% Bromophenol Blue) was diluted to 1X in the samples. The samples were then heated to 95ºC for 5 minutes and then loaded onto a Bio-Rad 4-20% Gradient Gel (TGX 10-Well catalog# 4561094, TGX 15-Well catalog# 4561096, Criterion TGX 18-well catalog# 5671094). The gels were run at 160V until the dye front ran off. The gels were then imaged for fluorescence at a wavelength of 535 nm on the Cytiva ImageQuant800 (catalog #29399484).

**Streptavidin Pulldown**

The Streptavidin Magnetic Beads (New England Biolabs, catalog #S1420S) were prepared by washing three times with a conjugation buffer (PBS + 500 mM NaCl), maintaining the beads in a 1:1 slurry. Per each reaction, the biotinylated sample was then mixed with 1 mL of conjugation buffer and 100 µL of the washed 1:1 Streptavidin bead slurry. The samples were then rotated for two hours at room temperature.

After rotating, the samples were washed nine times in total: three times with a wash buffer (PBS + 500 mM NaCl + 0.1% Tween 20), three times with PBS, and three times with MilliQ water.

**Flag-Immunoprecipitation**

Flag-tagged cells for MEK1, BRAF, and KSR2 from A375 cells were lysed resuspending cell pellets in 500 µL of a Lysis Buffer (50 mM Tris-HCl (pH 7.4–7.6 at 4°C), 140 mM NaCl, 1 mM EDTA, 1% NP-40, 0.2% Triton X-100, Protease inhibitor cocktail (Thermo Scientific cat#:PI78441)). Samples were then spun down at 16,000 g for 15 minutes at 4ºC. 25 µL of Sigma Flag-M2 affinity beads (50% slurry, cat#: A2220) were prepared per mg of protein. Beads were prepared by washing three times with a Wash Buffer (50 mM Tris-HCl (pH 7.4–7.6 at 4°C), 140 mM NaCl, 1 mM EDTA, Protease Inhibitor Cocktail), centrifuging the beads at 1,000 rpm for 1 minute. Bead were then equilibrated in Wash Buffer for 5 minutes on ice. Lysates were then added to equilibrated bead (4mg Flag-MEK1, 6mg Flag-KSR2, 9mg Flag-BRAFV600E), volumes were standardized to 1mL and samples were incubated on bead while rotating at 4°C overnight. After incubation, the beads were washed 4 times with 500 µL of Wash Buffer, centrifuging at 1,000 rpm for 1 minute between washes. Elution from the Flag-beads was performed by incubating the beads while rotating at 4°C for 4 hours with enough 100 µg/mL of 3x Flag Peptide (Sigma cat#:F4799) to match packed-bead volume.

**Co-Immunoprecipitation of BRAF**

BRAF Co-IP was performed with 1 mg of A375 lysate using Protein G Dynabeads (Thermo, cat#: 10004D) according to the manufacturers protocol, with the antibodies shown in Table S2.

**Immunoblot analysis**

The 250 µg samples were eluted off bead by adding 50 µL of a 5X Sample buffer and heating for 5 minutes in a water bath at 95ºC. After elution the samples were diluted so that the final concentration of the sample buffer was 1X, and a Magnetic Rack (New England Biolabs, catalog #S1509S) was used to separate out the beads. The supernatant was then run on a Bio-Rad 4-20% Gradient Gel.

For whole cell lysate, equal amounts of lysates were resolved by SDS-PAGE using standard techniques.

For all blots, the Transblot Turbo System (Bio-Rad, catalog #1704150EDU) was used to transfer the SDS-PAGE to PVDF using Trans-Blot Turbo RTA Midi 0.2 µm PVDF Transfer Kit (Bio-Rad, catalog #1704273) containing the Bio-Rad 5X Transfer Buffer (catalog #10026938).

Protein was detected with the primary antibodies (summarized in Table S2), followed by detection with one of the horseradish peroxidase conjugated secondary antibodies: goat anti-rabbit IgG (1:5000; Cell Signaling Technologies (CST), 7074) or goat anti-mouse IgG (1:5000; CST, 7076), using SignalFire (CST) or SignalFire Elite ECL (CST) detection reagents.

**Table S2: Primary Antibodies Used**

| Antibody | UniProt Accession IDs | Source (Cat#) | Dilution |
|---|---|---|---|
| Biotin | N/A | Cell Signaling (5597) | 1:5000 |
| β-Actin | P60709 | Cell Signaling (8457S) | 1:1000 |
| BRAF | P15056 | Cell Signaling (14814) | 1:1000 |
| phospho-BRAF (Ser445) | | Cell Signaling (2696) | 1:1000 |
| MEK 1/2 | Q02750, P36507 | Cell Signaling (4694) | 1:1000 |
| phospho-MEK 1/2 (Ser217/221) | | Cell Signaling (9154) | 1:1000 |
| ERK 1/2 | P27361, P28482 | Cell Signaling (4696) | 1:1000 |
| phospho-ERK1/2 (Thr303/Tyr204) | | Cell Signaling (4370) | 1:1000 |
| KSR1 | Q8IVT5 | Cell Signaling (4640) | 1:1000 |
| phospho-KSR1 (Ser392) | | Cell Signaling (4951) | 1:1000 |
| KSR2 | Q6VAB6 | ProSci (55-850) | 1:1000 |
| phospho-KSR2 (Thr497) | | Affinity Biosciences (AF4342) | 1:1000 |

| | | | |
|---|---|---|---|
| Flag | N/A | Cell Signaling (86861) | 1:2000 |

**Tryptic Digest**

On-bead tryptic digest was performed on the 1 mg samples. Fresh 100 mM DTT (Thermo Scientific, catalog #R0861) was added so that the final concentration in the samples was 10 mM, and the samples were incubated at 60°C for 20 minutes. Then fresh 10mM Iodoacetamide (Thermo Scientific, catalog #122271000) was added to each sample to a final concentration of 20 mM, and the samples were incubated at room temperature in the dark for 10 minutes. Finally, Trypsin/Lys C mix, Mass Spec Grade (Promega, catalog #V5072) was added to each sample in a protein ratio of 1:100. The samples were then incubated with shaking at 37°C overnight.

The next day, formic acid was added (0.1% of sample volume) to quench the reaction.

In-solution tryptic digest of flag-enriched MEK1, $BRAF^{V600E}$, and KSR2, and purified MEK1 and $BRAF^{V600E}$ were digested using this same protocol.

**Tandem mass tag (TMT) labeling**

Tandem mass tag (TMT) labeling of peptides was performed according to the manufacturer's instructions. Briefly, peptides underwent $C_{18}$ cleanup and were reconstituted in 100 mM triethylammonium bicarbonate (TEABC) buffer (pH ~8.0). The TMT kit was equilibrated to room temperature, and each tag was reconstituted in 50 µL of anhydrous acetonitrile (ACN) with gentle vortexing. Three independent TMT experiments were set up: one with the cysteine reactive probe IA-alkyne, another with the lysine reactive probe ArSq-alkyne, and one without any probe. In each of these probe conditions, there were three replicates of A375 samples treated with vehicle (DMSO), 100 nM Trametinib, and 1 µM Vemurafenib. Therefore, nine of the 50 µL aliquots of each label from the TMT sixteen-plex kit (TMT126-130C, ThermoFisher, catalog#A44520) were resuspended, and one tag each was added to each of the samples in each probe condition.

TMT reagents were added to the corresponding samples as detailed in Table S3, and the labeling reactions were incubated at room temperature for 1 hour. To assess labeling efficiency, an equal volume (2 µL) from each sample was pooled and labelling efficiency determined by LC-MS/MS. Following the label check evaluation, the reaction was quenched by adding 5% hydroxylamine and incubated for 15 minutes. Samples were then pooled and vacuum dried.

To enhance proteome depth, $C_{18}$ spin column-based fractionation was performed using a stepwise acetonitrile gradient (5–50%). Each sample was fractionated into 14 fractions, which were subsequently concatenated into 7 fractions. Samples were vacuum dried and stored at -20°C until LC-MS/MS analysis.

**Table S3: Experimental sample details and TMT labeling layout for quantitative proteomic analysis.**

| Sample Name | Replicate Number | TMT label used |
|---|---|---|
| IA-Alkyne | R1 | 126 |
| IA-Alkyne | R2 | 127N |
| IA-Alkyne | R3 | 127C |
| IA-Alkyne (Trametinib) | R1 | 128N |
| IA-Alkyne (Trametinib) | R2 | 128C |
| IA-Alkyne (Trametinib) | R3 | 129N |
| IA-Alkyne (Vemurafenib) | R1 | 129C |
| IA-Alkyne (Vemurafenib) | R2 | 130N |
| IA-Alkyne (Vemurafenib) | R3 | 130C |
| ArSq-alkyne | R1 | 126 |
| ArSq-alkyne | R2 | 127N |
| ArSq-alkyne | R3 | 127C |
| ArSq-alkyne (Trametinib) | R1 | 128N |
| ArSq-alkyne (Trametinib) | R2 | 128C |
| ArSq-alkyne (Trametinib) | R3 | 129N |
| ArSq-alkyne (Vemurafenib) | R1 | 129C |
| ArSq-alkyne (Vemurafenib) | R2 | 130N |
| ArSq-alkyne (Vemurafenib) | R3 | 130C |
| No Probe, No inhibitor | R1 | 126 |
| No Probe, No inhibitor | R2 | 127N |
| No Probe, No inhibitor | R3 | 127C |
| No Probe (Trametinib) | R1 | 128N |
| No Probe (Trametinib) | R2 | 128C |
| No Probe (Trametinib) | R3 | 129N |
| No Probe (Vemurafenib) | R1 | 129C |
| No Probe (Vemurafenib) | R2 | 130N |
| No Probe (Vemurafenib) | R3 | 130C |

**Orbitrap LC-MS/MS Analysis**

LC-MS/MS analysis was performed on a Q Exactive Hybrid Quadrupole-Orbitrap mass spectrometer (Thermo Scientific, Bremen, Germany) coupled with a Vanquish Neo UHPLC system (Thermo Scientific). Labelled peptides were reconstituted in 0.1% formic acid and separated on an analytical column (75 µm × 15 cm; 3 µm $C_{18}$ 100Å; Thermo Scientific, PepMap™ RSLC, PN ES900) at a flow rate of 350 nL/min. A linear gradient of 4–35% solvent B (0.1% formic acid in 80% acetonitrile) was applied over 100 minutes, with a total run time of 120 minutes.

MS and MS/MS data were acquired using the Orbitrap analyzer at resolutions of 70,000 and 35,000, respectively. Precursor MS scans were performed in the m/z range of 350–1500, with a maximum ion injection time of 50 msec and an AGC target of $3e^6$ ions. The mass spectrometer operated in DDA mode, selecting top 10 precursor ions using a quadrupole mass filter with an isolation window of 1.2 m/z and a dynamic exclusion of 45 seconds. Fragmentation of isolated ions was performed using high-energy collision-induced dissociation (HCD) with an NCE of 30%. MS/MS scans were acquired with a maximum ion injection time of 250 msec and an AGC target of $2e^5$.

#### Data Analysis

Data analysis was performed using the Proteome Discoverer platform (version 3.0.0.757; Thermo Scientific). Mass spectrometry raw (.raw) files were searched against the Human UniProt database using the SEQUEST search algorithm. TMT labeling at the peptide N-terminus/lysine residue were set as static modifications, while methionine oxidation and carbamidomethylation at cysteine residues was considered variable modification. False discovery rate (FDR) of 5% was applied at both the PSM and peptide levels and a precursor mass tolerance of 20 ppm and a fragment mass tolerance of 0.05 Da were used. Trypsin was specified as the protease, allowing a maximum of two missed cleavages (three missed cleavages were allowed for ArSq labelled proteins).

#### QTOF LC-MS/MS Analysis

LC/MS analyses were conducted on an Agilent 1290 Infinity II UHPLC system coupled with an Agilent 6545XT AdvanceBio LC/Q-TOF system equipped with an Agilent Dual Jet Stream ESI source. LC separation was obtained with an Agilent AdvanceBio Peptide Mapping column (2.1 × 150 mm, 2.7 µm). Peptides were separated over a gradient of 1-95% acetonitrile in water (both containing 0.1% formic acid) over 45 minutes with a flow rate of 0.4 ml/min. MS and MS/MS data were acquired using the Q-TOF with the following parameters:

| | |
|---|---|
| Drying Gas | 11 L/min |
| Drying Gas Temperature | 325 °C |
| Sheath Gas Flow | 10 L/min |
| Sheath Gas Temperature | 325 °C |
| Nebulizer Pressure | 35 psi |
| Capillary Voltage | 4,000 V |
| Nozzle Voltage | 0 V |
| Fragmentor Voltage | 175 V |
| Acquisition Mode | AutoMS2 |

#### Data Analysis

Data analysis was performed using Agilent MassHunter BioConfirm. Mass spectrometry files were searched against the relevant protein sequences and the noted chemical adducts were included as variable modifications, as well as carbamidomethylation at cysteine and methionine oxidation. Trypsin was specified as the protease, allowing a maximum of two missed cleavages (three missed cleavages were allowed for ArSq labelled proteins). A precursor mass tolerance of 5 ppm and a fragment mass tolerance of 20 ppm Da were used.

#### Additional Supplementary Files

Table S4 - IA-Alkyne Competition Proteomics

Table S5 - ArSq-Alkyne Competition Proteomics

Table S6 – Identified Kinases from IA-alkyne and ArSq-alkyne Competition Proteomics
